# Sequence Analysis and Homology Modelling of SmHQT Protein, a Key Player in Chlorogenic Acid Pathway of Eggplant

**DOI:** 10.1101/599282

**Authors:** Prashant Kaushik, Dinesh Kumar Saini

## Abstract

Eggplant is an important vegetable that belongs to family Solanaceae. Fruits of eggplant are rich in phenolic acids. Chlorogenic acid makes up to 90 per cent of total phenolic acids present in the eggplants fruit flesh. Eggplant hydroxycinnamoyl CoA-quinate (SmHQT) is the central enzyme that modulates the last step of the chlorogenic acid pathway of eggplant. Here, we have analysed the sequence of eggplant SmHQT protein in eggplant. The sequence obtained from the NCBI was aligned using MUSCLE. After that, homology modelling was performed using MODELLER 9v15. Model with Dope Z-Score of −1.596 was selected and verified for viability under real conditions using several online tools. Also, the docking was performed with this model. Overall, this could be useful in developing eggplant varieties rich in phenolic acids especially chlorogenic acid.

## Introduction

Vegetables produce several bioactive compounds like phenolic acids (Kaushik *et al.*, 2015; Goleniowski *et al.*, 2013; Harborne, 1984). Eggplant (*Solanum melongena* L.) encompasses a high concentration of phenolic acids, which are beneficial for human health and development (Kaushik *et al.*, 2017; Rice-Evans *et al.*, 1996; Robbins, 2003). Within the eggplant flesh, the chlorogenic acid exists as an ester as 5-caffeoylquinic acid, and it tends to make as much as 90 *%* of total phenolics within the eggplant flesh (Meyer *et al.*, 2015; Stommel and Whitaker, 2003). Chlorogenic acid biosynthesis pathway in eggplant was mapped. The hydroxycinnamoyl CoA-quinate (HQT) is the key enzyme tested to boost the chlorogenic acid content in Solanaceae (Tomato and Potato) as well as in other related families (Niggeweg *et al.*, 2004). Although there is not any detailed information regarding the SmHQT, therefore, here firstly we have analysed the amino acid sequences present in the SmHQT of eggplant. After that, using homology modelling methods 3D model of SmHQT was predicated. Subsequently, several properties of modelled structure like stereochemical properties etc. were studied.

## Material and Methods

### Protein sequence analyses

The SmHQT of *S. melongena* (NCBI protein accession number: AMK01803) is 427 amino acids long. First of all, the ProtParam tool (http://web.expasy.org/protparam/) (Gasteiger *et al.*, 2005) of ExPASy was used to compute the following physicochemical properties: molecular weight, theoretical isoelectric point (pI), amino acid composition (%), number of total negatively and positively charged residues, aliphatic index, instability index, and Grand Average of Hydropathy (GRAVY). For this, the SWISS-PROT ID of protein (ID: A0A126Q9A5) was used. Similarly, properties at the secondary level were predicted using an online tool PredictProtein (https://www.predictprotein.org/) (Yachdav *et al.*, 2014). Using MEME suite (http://meme-suite.org/) (Timothy *et al.*, 1994), motif search was performed with following parameters: some repetitions being any; a maximum number of motifs being 6; optimum width of the motif being 50, and the site being 2-600. iPSORT (http://ipsort.hgc.jp/) (Bannai *et al.*, 2002) tool was used to predict N-terminal sorting signals, and ProtComp V9.0 program (http://www.softberry.com/berry.phtml) was employed for the identification of sub-cellular localisation of the protein.

### Gene ontology

Gene ontology terms were predicted by an online tool MetaStudent (https://bio.tools/metastudent) (Hamp *et al.*, 2013). These terms were organized in three different ontologies, molecular function ontology, biological process ontology and cellular component ontology. Each predicted term was associated based on a reliability score (0–100), indicating the chance that the protein has that function.

### Prediction of transmembrane helices

TMHMM Server v. 2.0 (Krogh *et al.*, 2001) was employed to predict transmembrane helices in SmHQT protein of *S. melongena.*

### Protein-RNA association

catRAPID algorithm (http://s.tartaglialab.com/page/catrapid_group) (Agostini *et al.*, 2013) was used to estimate the binding propensity of protein-RNA pairs. Where a prediction score of less than 0.5 suggests non-RNA binding nature of the protein. For all the above analysis, the amino acid sequence of protein was uploaded.

### Sequence alignment and homology modelling of SmHQT

The SmHQT protein sequence of *Solanum melongena* (NCBI accession number: AMK01803) was retrieved from NCBI (https://www.ncbi.nlm.nih.gov). Homology modelling SmHQT of *Solanum melongena* was performed in the following steps: template selection using online tools HHblits and BLASTp, sequence-template alignment, model building and validation.

### Template selection, sequence-template alignment

The sequence homology search was carried out by using the online tools HHblits (https://toolkit.tuebingen.mpg.de/#/tools/hhblits) (Remmert *et al.*, 2012) and BLASTp. Out of the total hits, two best templates were selected based on coverage, resolution and identity percentage. The amino acid sequences of two selected proteins were retrieved from NCBI and aligned using MUSCLE alignment tool (https://www.ebi.ac.uk/Tools/msa/muscle/) (Edgar, 2004). The aligned sequence file (Fig. 1) was used as a template to generate a comparative 3 D model of SmHQT of *Solanum melongena* by MODELLER 9v15 (Eswar *et al.*, 2008). MODELLER 9v15 builds an atomic model based on an experimentally determined structure that is closely related at the sequence level.

**Fig. 1:**
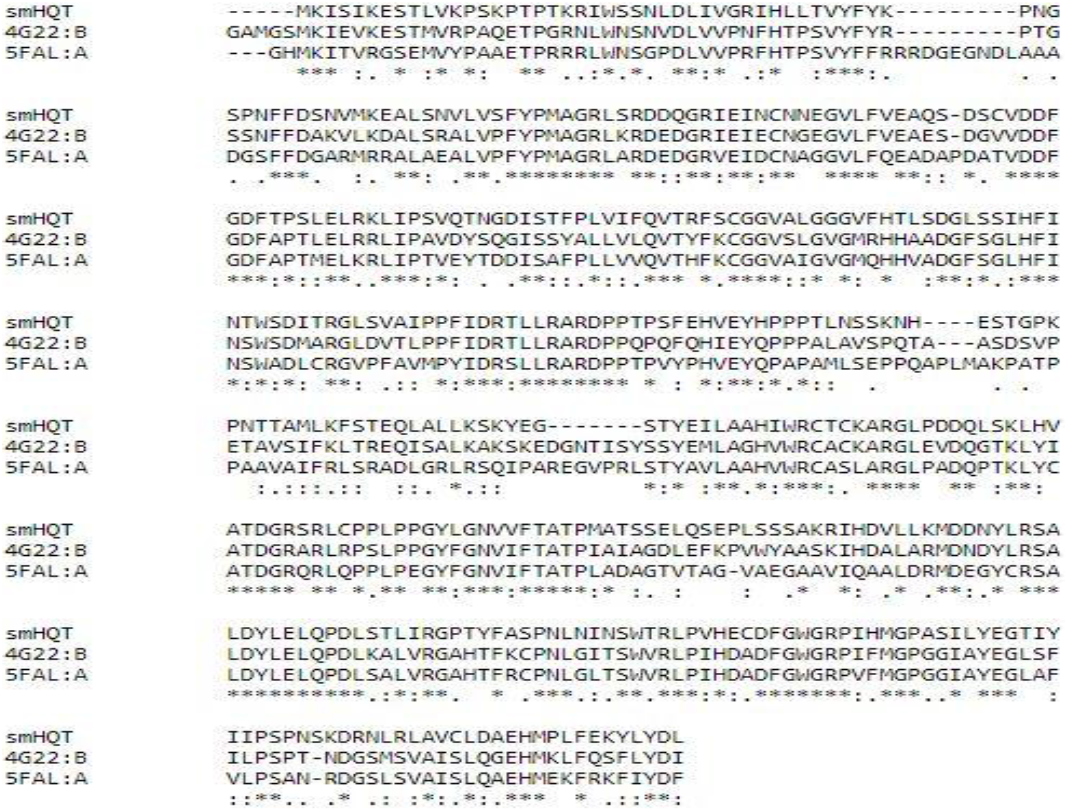
Sequence alignment of SmHQT of *S. melongena* and 4G22_B and 5FAL_A (Templates).

### Model building

MODELLER 9v15 generated several preliminary models which were ranked based on their discrete optimised protein energy (DOPE) scores (Table 1). Model (model002) having the lowest DOPE score was selected and further subjected to quality assessment.

**Table 1:**
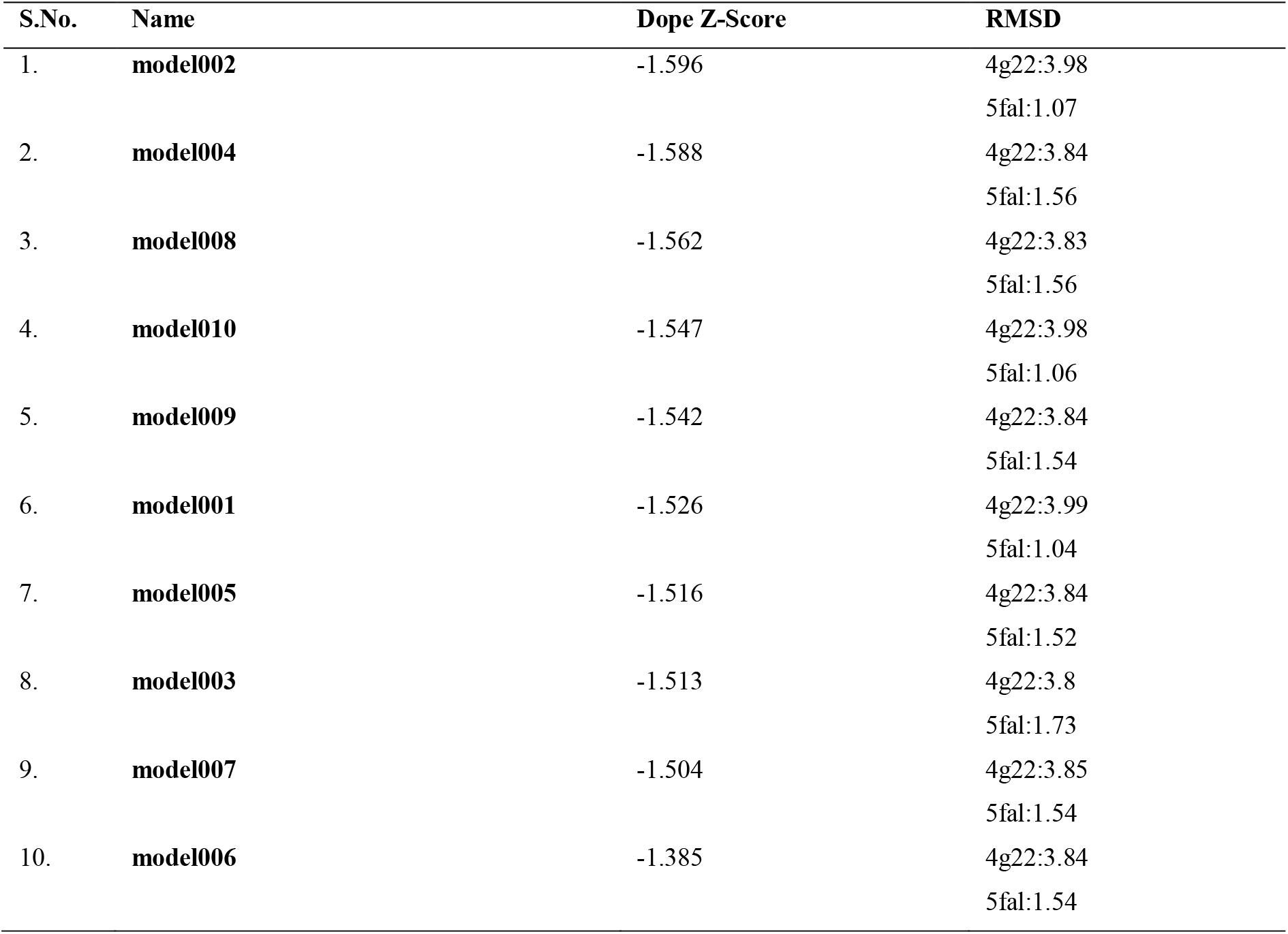
Model002 is selected based on Dope Z-Score.

### Verification of the generated 3D structure model

The PROCHECK server (http://servicesn.mbi.ucla.edu/PROCHECK/) (Laskowski, 1993) was used for evaluating the quality of the modelled SmHQT *SmHQT*protein structure concerning energy and stereo-chemical geometry. Ramachandran plot was prepared to find out the relative proportion of amino acids (aa), which fall in the favoured region, relative to other regions (Laskowski, 1993). QMEAN4 (Quantitative Model Energy Analysis) (Benkert *et al.*, 2010) which is a linear combination of four potential statistical terms, was used to derive both global (i.e. for the entire structure) and local (i.e. per residue) absolute quality estimates by one single model. The QMEAN Z-score was used for measuring the absolute quality of the model. QMEANDisCo (Waterhouse *et al.*, 2018) was used to improve the local QMEAN quality estimates with a new term directly derived from structures of homologous proteins. Using ERRAT program (Colovos and Yeates., 1993) from SAVES (Structure Analysis and Verification Server), non-bonded interactions were analysed, and the quality factor for modelled protein was recorded in percentage. Compatibility of an atomic model (3D) with its amino acid sequence (1D) was determined with the help of VERIFY-3D program (Eisenberg *et al.*, 1997) from SAVES and the results were obtained (3D-1D Profile) in percentage.

The ProSA (Wiederstein and Sippl, 2007) web server was employed to evaluate the energy of the model protein related to good protein structure. PROSESS (Protein Structure Evaluation Suite & Server) web server (http://www.prosess.ca/) was also employed for evaluating and validating the modelled protein structure (Berjanskii *et al.*, 2010). For further assessment of modelled protein at a global and local level, eQuant (https://biosciences.hs-mittweida.de/equant/) was also used (Bittrich *et al.*, 2015). It expressed the overall quality using the global distance test (GDTscore), which was used to quantify the predicted degree of structural deviation between the processed structure model and the unknown native structure, i.e. modelled protein. It also predicted the distances between corresponding residues which were further used for predicting the local quality of model.

### Functional annotation

PROFUNC server (Laskowski *et al.*, 2005) server was used for predicting the protein function from 3D structure.

### Identification of secondary structure in modelled protein

STRIDE (Structural identification) algorithm (http://webclu.bio.wzw.tum.de/cgibin/stride/stridecgi.py) (Heinig and Frishman, 2004) was used for assigning the protein secondary structure elements in the atomic coordinates of the protein. For this purpose, model file was uploaded.

### Structure-based protein stability prediction

MAESTRO web server (https://biwww.che.sbg.ac.at/maestro/web) (Laimer *et al.*, 2016) was used for three different tasks viz. to calculate a mutation sensitivity profile, to perform a search for the most (de) stabilizing n-point mutations and to evaluate potential disulphide bonds. Mutation sensitivity profile shows the impact of each mutation at each possible position which allows the identification of cold or hot spot sites of mutations. ΔΔGpred. (in kcal/mol), Sss and Cpred. values indicating total predicted changes in stability, bond score and confidence estimate respectively, were calculated. For all above tasks, model file was uploaded.

### Detection of protein-protein interaction sites

BindML/BindML+ web servers (http://kiharalab.org/bindml/plus/) (Wei *et al.*, 2017) were used for predicting the protein-protein interaction sites by identifying mutation patterns found in known protein-protein complexes, and for distinguishing the permanent and transient types of protein-protein interaction sites, respectively. For this purpose, we uploaded PDB file of model protein. We left MSA filed empty and allowed the server to execute a search against the Pfam database and to generate the MSA file with the MUSCLE multiple sequence alignment program.

### Molecular docking

The SwissDock (http://www.swissdock.ch/docking) program (Grosdidier *et al.*, 2011) was used for the molecular docking analysis of modelled SmHQT protein with substrate, i.e. quinic acid. The two-dimensional structure of substrate quinic acid was retrieved from the pubchem (Wang *et al.*, 2009) server of NCBI and then transformed into the 3D coordinate via the LINUX using AMBER tool. This 3D coordinate was then docked against modelled protein using the mentioned software. The target molecule was provided as a PDB file. The docking process from SwissDock was set to the accurate type and definition of the interest was set as default. The allowance for the flexibility for side chains was set to within 0 Å of any atom of the ligand in its reference binding mode.

## Results

### Protein sequence analyses

The molecular weight of SmHQT protein sequence was 47.55KD. The total number of negatively charged aa residues (Asp + Glu =44) exceeded the total number of positively charged aa residues (Arg + Lys =41). The isoelectric point was within the acidic range (6.46). SmHQT protein was found to be unstable (instability index=45.28). The aliphatic index and GRAVY were 88.13 and −0.204 respectively. Predicted secondary structure of the protein was comprised of 23.4%, 22.2% and 54.3%; helix, strand and loops respectively. This protein was not found to contain any nuclear localization signal. Six distinct motifs were present in SmHQT proteins as identified by MEME suite. The details regarding their E-value, site count and width are given (Table 2). Using iPSORT online tool, a mitochondrial targeting peptide was found at the N-terminus of the smQHT protein. Further analysis through ProtComp predicted SmHQT protein to be localized in the mitochondria.

**Table 2:** Details related to motifs.

| S.No. | E-value | Sites | Width |
| --- | --- | --- | --- |
| 1. | 1.5e+001 | 2 | 7 |
| 2. | 2.1e+001 | 2 | 8 |
| 3. | 5.4e+001 | 2 | 6 |
| 4. | 3.1e+001 | 2 | 6 |
| 5. | 5.5e+001 | 2 | 6 |
| 6. | 2.7e+001 | 2 | 7 |

### Gene ontology

Gene ontology (GO) analysis suggested that SmHQT protein has transferase activity with their roles in transferring acyl groups other than amino-acyl groups and involves in metabolic processes. All computed GO-ID, GO-terms with their reliability *%* are given (Supplementary Fig. 1,2,3).

### Prediction of transmembrane helices and Protein-RNA association

No transmembrane helix in *SmHQT* protein of *S. melongna* was predicted using TMHMM Server v. 2.0. We obtained a prediction score less than 0.5 (= 0.32) using the catRAPID algorithm which suggests no binding of SmHQT protein with RNA.

### Sequence alignment and homology modelling of *hydroxycinnamoyl coenzyme A-quinate transferase* (*SmHQT*)

The protein sequence of SmHQT was used for sequence homology search using the tools HHblits and BLASTp with default parameters. Out of the total hits, two proteins viz. hydroxycinnamoyl-CoA shikimate/quinate hydroxyl cinnamoyl transferase from *Coffea canephora* (PDB ID: 4G22_B) and hydroxycinnamoyl-CoA shikimate/quinate hydroxyl cinnamoyl transferase (PDB ID: 5FAL_A) were selected on the basis of coverage (100%), lowest e-values (7.6e-44 and 7.2e-43), good resolutions (1.7 Å and 1.861 Å), good identities (57 % and 53%) and then used as templates for homology modelling. An alignment of SmHQT protein sequence of *S. melongena* with the template protein sequences of hydroxycinnamoyl-CoA shikimate/quinate hydroxyl cinnamoyl transferase of *Coffea canephora* (PDB ID: 4G22_B) and hydroxycinnamoyl-CoA shikimate/quinate hydroxyl cinnamoyl transferase (PDB ID: 5FAL_A) of *Panicum virgatum* is shown in Fig. 1. MODELLER 9v15 was used to generate the homology model of SmHQT protein of *S. melogena* according to the crystal structures of 4G22_B and 5FAL_A. A total of ten models were generated and their discrete optimized potential energy (DOPE) scores were recorded (Table 1). The model002 having maximum score was considered as the best model of SmHQT of *S. melongena.* To visualize the model, a molecular visualisation system, PyMOL (DeLano, 2002) software was used (Fig. 3).

**Fig. 2:**
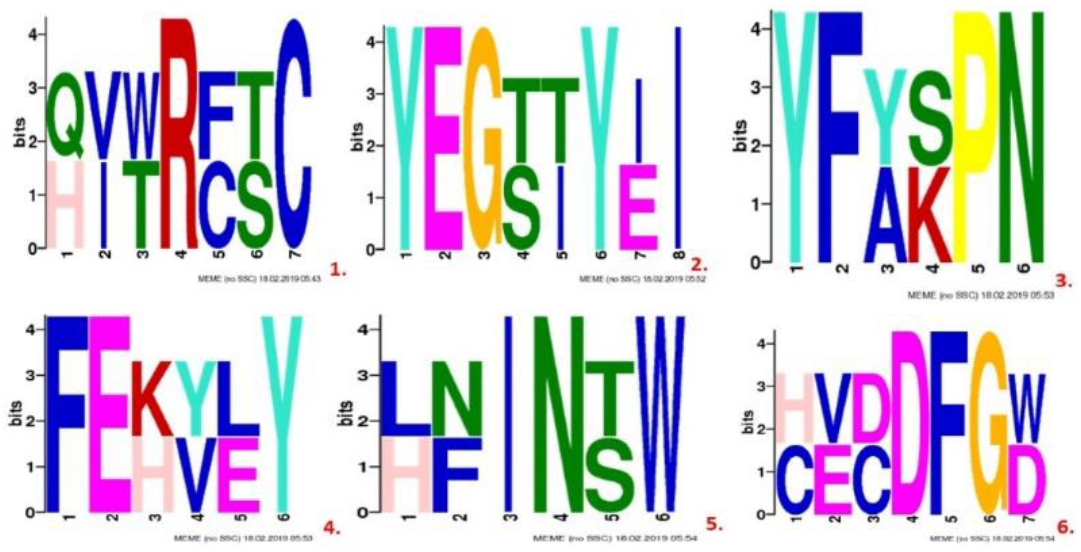
Representative figures of motifs.

**Fig. 3:**
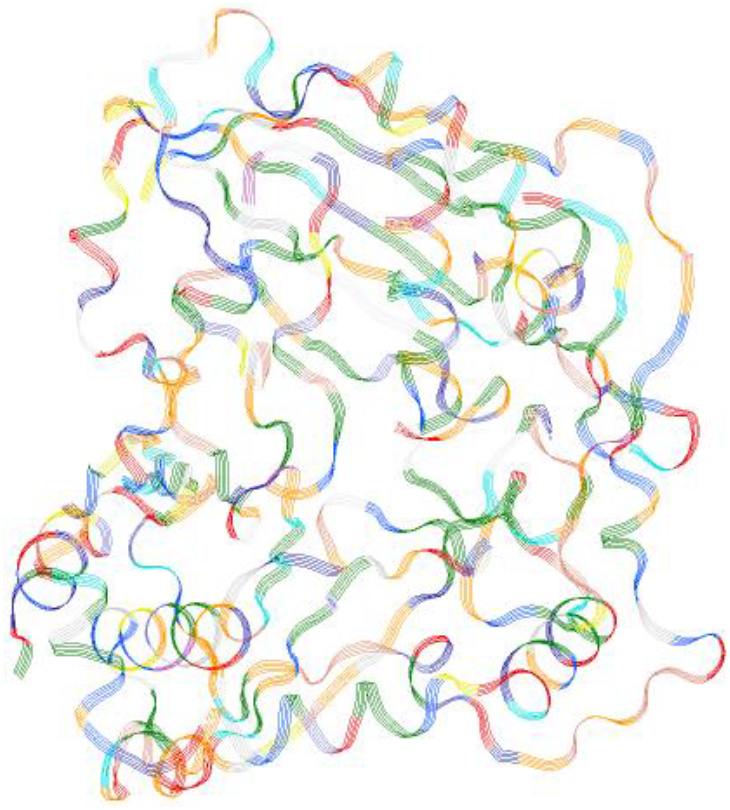
Visualization of 3D structure model002

### Verification of generated 3D structure model

PROCHECK, VERIFY3D and ERRAT programs were used for validating the predicted model. PROCHECK analysis provided a plot named as Ramachandran plot of the modelled protein showing distribution patterns of residues in different regions (Fig. 4). Among the 427 residues, 340 residues (93.2%) were found in the most favoured region, 24 (6.6%) in additional allowed region, one residue (0.3%) in the generously allowed region and no residue (0%) in the disallowed region (Table 4).

**Fig. 4:**
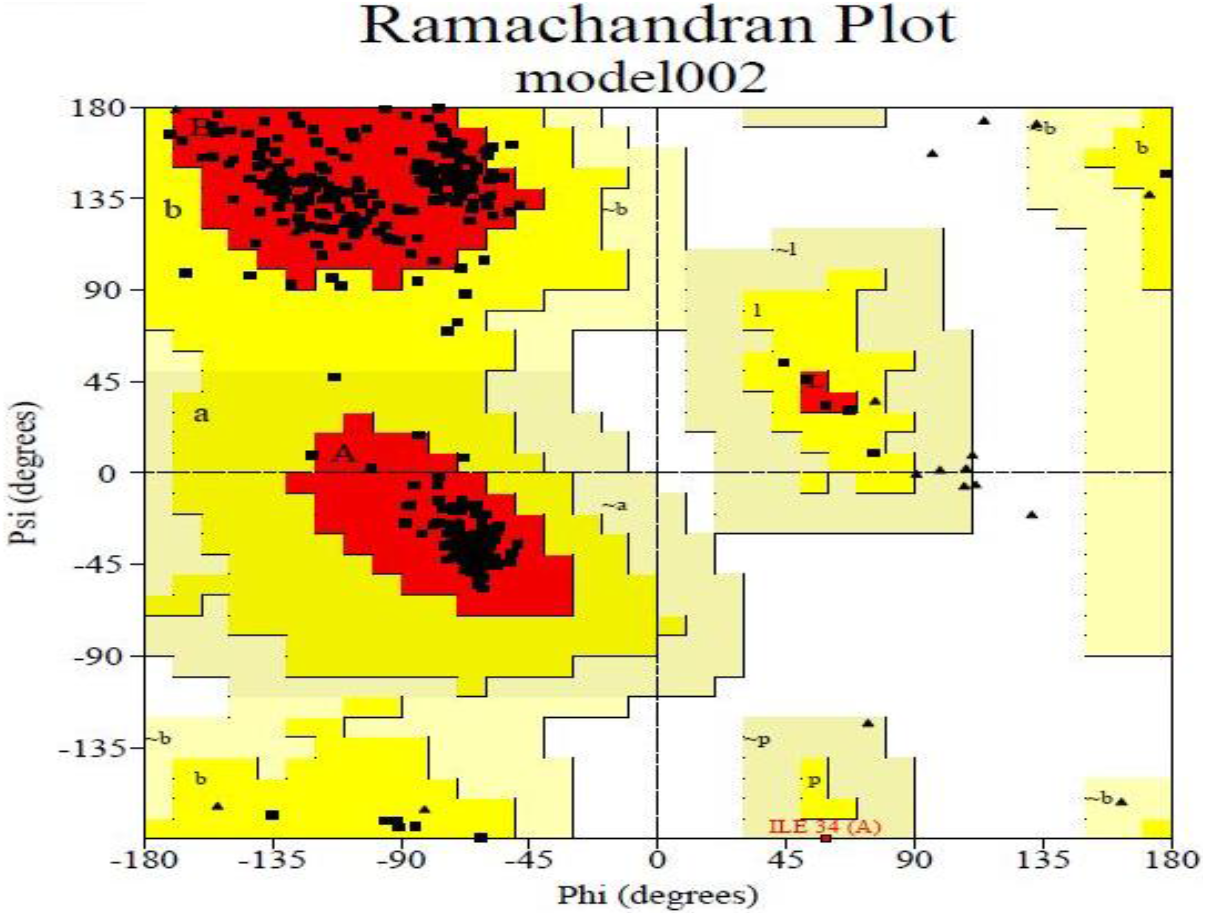
Ramachandran plot of modelled protein of *SmHQT.*

**Table 3:** Gene ontology (GO) terms with their reliability (%) to SmHQT via homology to already annotated proteins.

| GO ID | GO Term | Reliability |
| --- | --- | --- |
| Molecular Function Ontology |  |  |
| GO:0016747 | transferase activity | 39 |
| GO:0016740 | transferase activity | 38 |
| GO:0016746 | transferase activity, transferring acyl groups | 38 |
| GO:0003824 | catalytic activity | 27 |
| GO:0008374 | O-acyltransferase activity | 26 |
| GO:0050737 | O-hydroxycinnamoyltransferase activity | 26 |
| GO:0047172 | shikimate O-hydroxycinnamoyltransferase activity | 26 |
| GO:0050734 | hydroxycinnamoyltransferase activity | 25 |
| GO:0005488 | binding | 20 |
| GO:0050266 | rosmarinate synthase activity | 17 |
| Biological Process Ontology |  |  |
| GO:0008152 | metabolic process | 43 |
| GO:1901576 | organic substance biosynthetic process | 17 |
| GO:0071704 | organic substance metabolic process | 17 |
| GO:0009058 | biosynthetic process | 17 |
| GO:0044710 | single-organism metabolic process | 16 |
| GO:0006950 | response to stress | 16 |
| GO:0044249 | cellular biosynthetic process | 16 |
| GO:0044699 | single-organism process | 16 |
| GO:0050896 | response to stimulus | 16 |
| GO:0009987 | cellular process | 16 |
| Cellular Component Ontology |  |  |
| GO:0005737 | cytoplasm | 52 |
| GO:0044464 | cell part | 52 |
| GO:0005623 | cell | 52 |
| GO:0005622 | intracellular | 52 |
| GO:0044424 | intracellular part | 52 |
| GO:0005829 | cytosol | 38 |
| GO:0044444 | cytoplasmic part | 38 |
| GO:0043229 | intracellular organelle | 17 |
| GO:0043227 | membrane-bounded organelle | 17 |
| GO:0043226 | organelle | 17 |

**Table 4:**
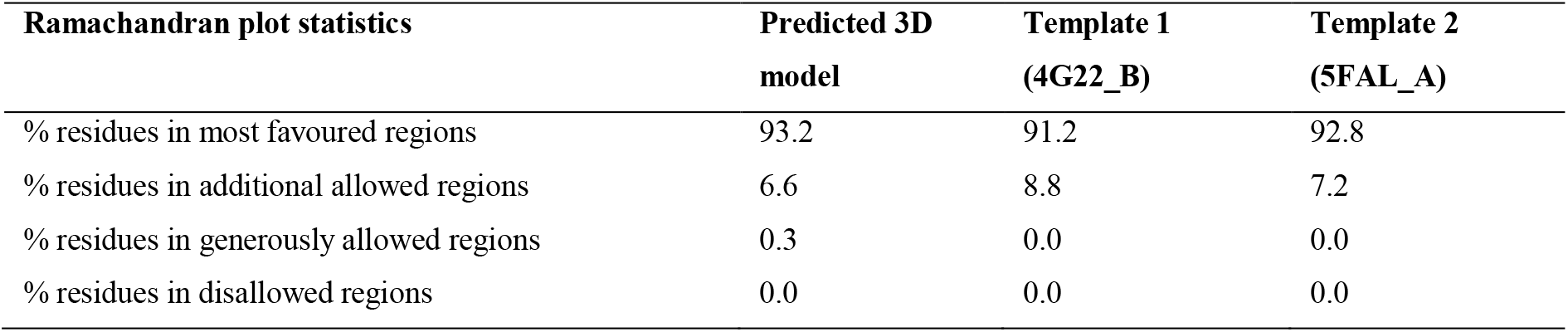
Comparing Ramachandran plot calculations of predicted 3D model with templates.

Ramachandran plots showing how each residue’s phi-psi dihedral angles vary for each residue across the different models in the NMR ensemble are also given (Supplementary Fig. 4). Dark shading on Chi1-Chi2 plots indicated favourable regions covered by residues on plots (Supplementary Fig. 5). Main chain parameters plot (Supplementary Fig. 6) showed six properties viz. Ramachandran plot quality, peptide bond planarity, bad non-bonded interactions, calpha tetrahedral distortion, main-chain hydrogen bond energy and overall G-factor. The values for all six parameters are also given (Supplementary Fig. 6) indicating the quality of the predicted model. Results obtained from PROCHECK indicate the reliability of the selected protein model.

A comparison of the predicted model with a non-redundant set of PDB structures (high-resolution X-ray structures of similar size) was also made (Fig. 5). On the graph, our model is represented by a red star. Local QMEAN score was also compared with local DisCo scores and the resulting QMEANDisCo scores (Fig. 6). The quality factor of our predicted model was 68.74. Protein model is suggested to be of good quality if at least 80% of the amino acids of the protein score >= 0.2 in the 3D/1D profile. In our case, 90.63% of the residues of model protein scored >= 0.2 in the 3D/1D profile (Fig. 7). Overall, the selected model appears to be of good quality, as inferred firstly from the acceptable values of QMEAN4 and secondly from the high values of quality factors estimated by ERRAT and those of 3D-1D score estimated by VERIFY-3D. The Z score of the protein represents the overall quality and measures the deviation of the total energy of the protein structure. The Z score of the protein is displayed in this plot with a dark black point (Fig. 8). Groups of structures determined from different sources (X-ray, NMR) are distinguished by different colours (X-ray in light blue and NMR in dark blue). In this plot, the Z score value of the predicted model (model002) of SmHQT of *S. melongena* (−9.57) is located within the space of protein related to X-ray. This Z-score of model protein is very close to the score of the templates (−10.35 and −10.71) which suggests that the predicted model is reliable and very close to the experimentally determined structure. A plot which shows local model quality by plotting energies as a function of amino acid sequence position is represented (Fig. 9).

**Fig. 5:**
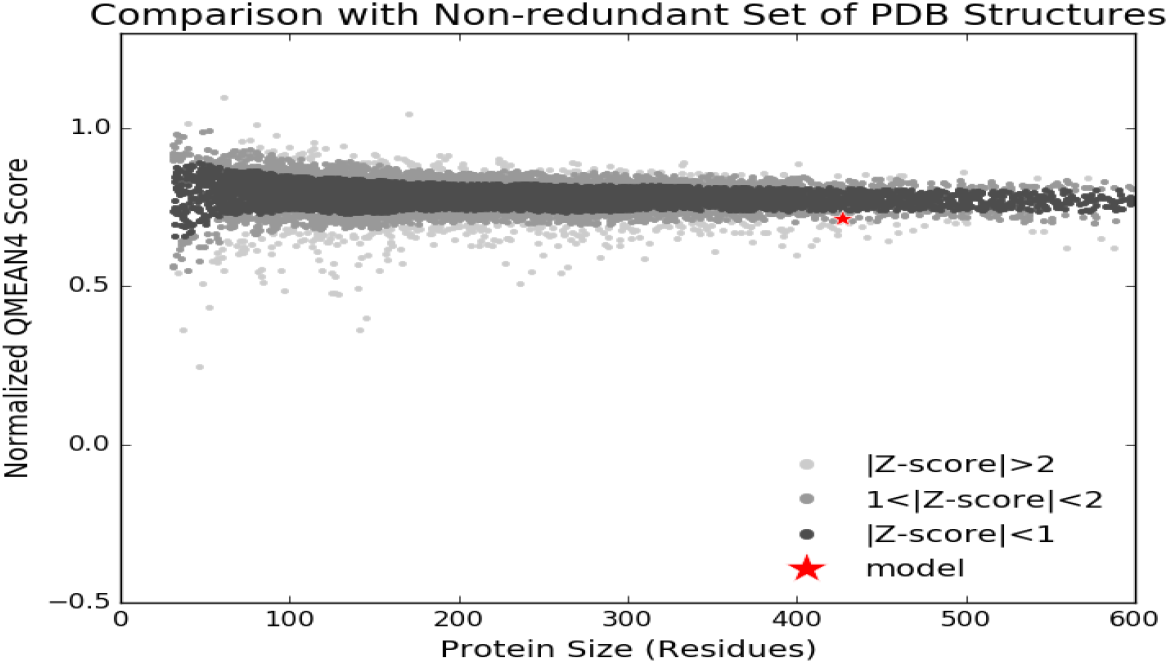
Comparison of predicted model with non-redundant set of PDB structures.

**Fig. 6:**
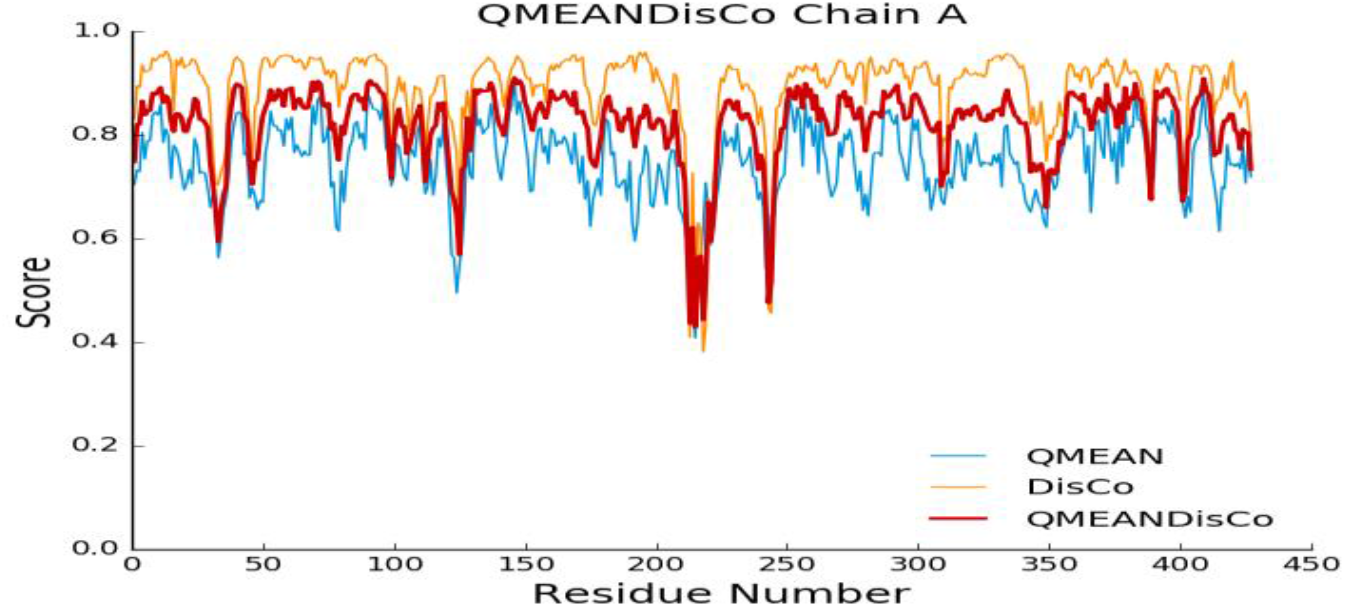
Comparison of local QMEAN score with local DisCo and QMEANDisCo scores.

**Fig. 7:**
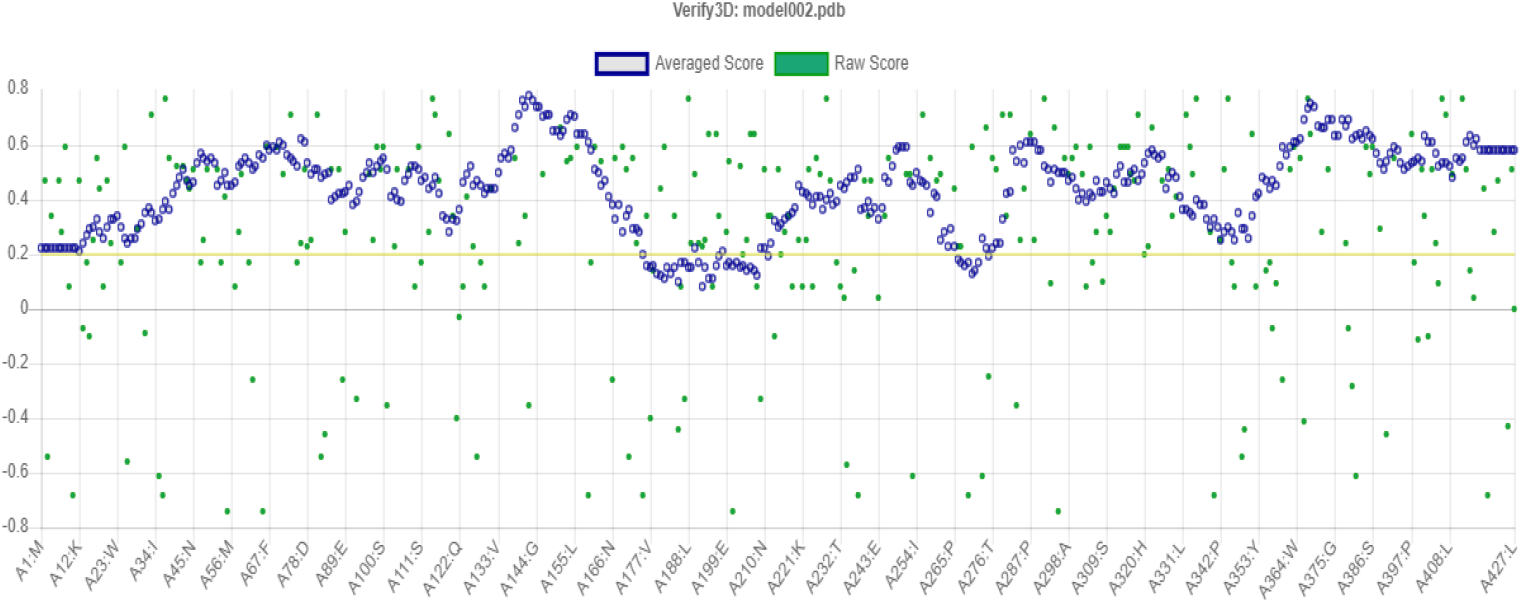
Verify 3D score profile.

**Fig. 8:**
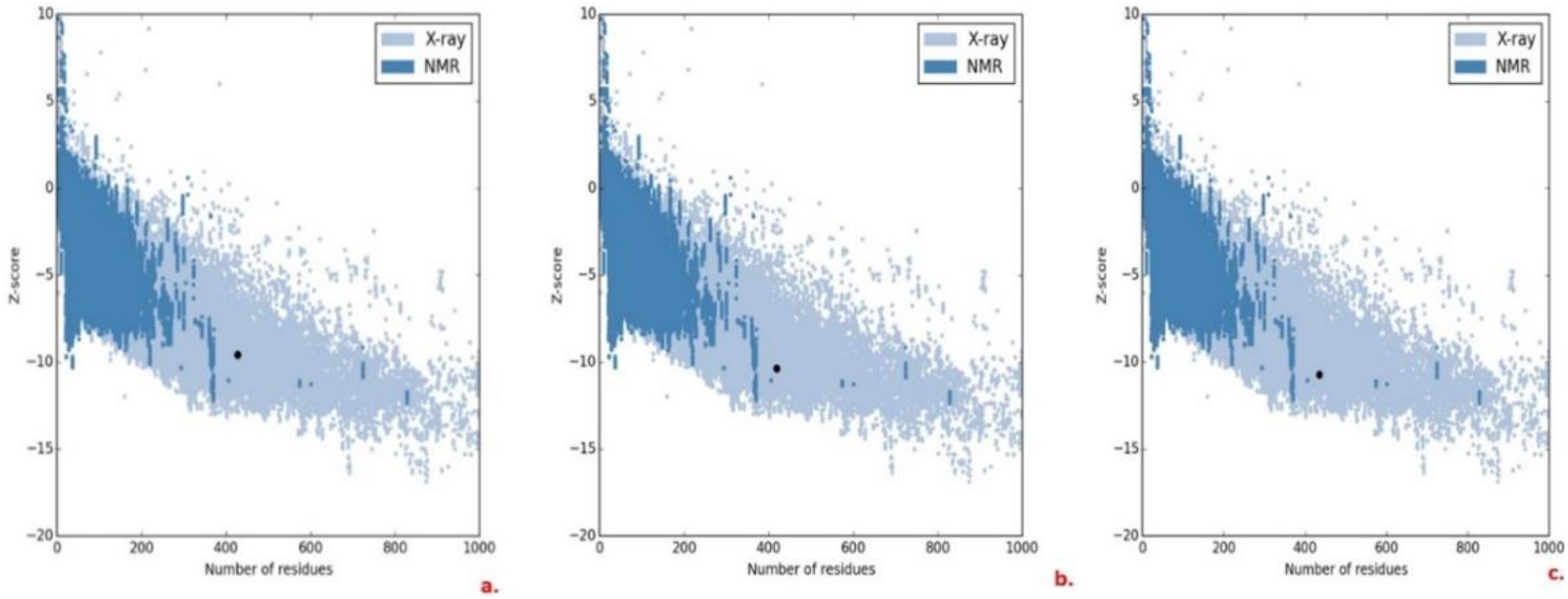
Z-Score plots showing Z scores of proteins (the three black dots represent Z-score of the model protein (a) and templates viz. (b) 4G22_B and (c) 5FAL_A).

**Fig. 9:**
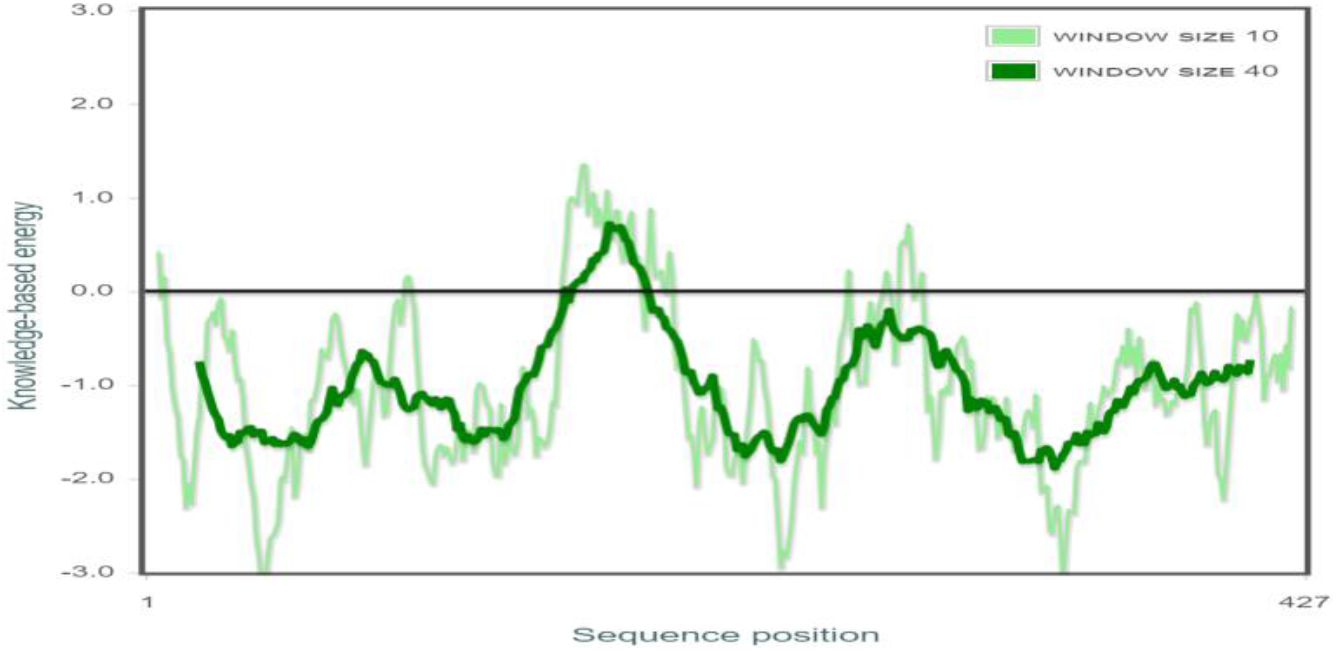
Energy plot for the predicted SmHQT protein by ProSA

Global structure assessment was also performed with PROSESS. Our predicted model achieved overall quality, covalent bond quality, non-covalent/packing quality and torsion angle quality within an acceptable range (Fig. 10). Assessment with eQuant predicted GDT, Z-score and TM-score as 78.58, 1.82 and 0.86 respectively. A score of 78.58 shows the good global quality of modelled protein (Fig. 11). Server predicted the distance between corresponding residues and plotted on the y-axis. Residues with distances exceeding 3.8 Å (shaded red) indicate local discrepancies (Fig. 12). STRIDE server revealed 116 (27.16%) residues in alpha-helix, 99 (23.18%) residues in β strands, 89 (20.84%) residues in coils, 100 (23.41%) residues in β turns, 21 (4.91%) in 310 helix and two (0.46%) in bridge (Fig. 13).

**Fig. 10:**
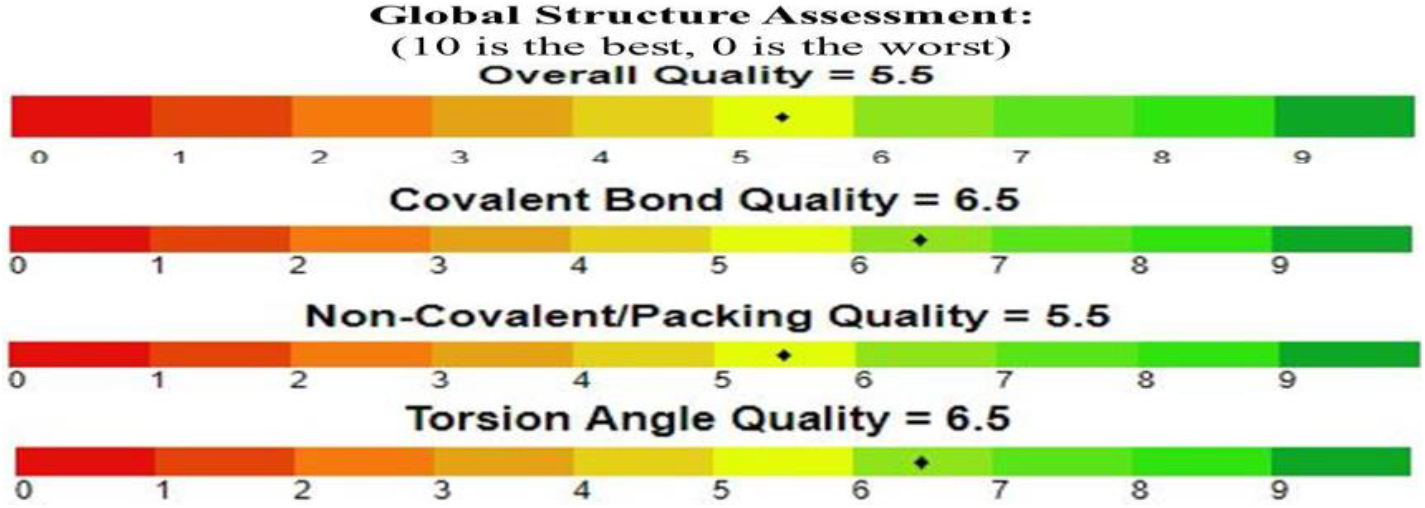
Showing different quality parameters of predicted model (represented by black star on scale) on a 0–10 scale.

**Fig. 11:**
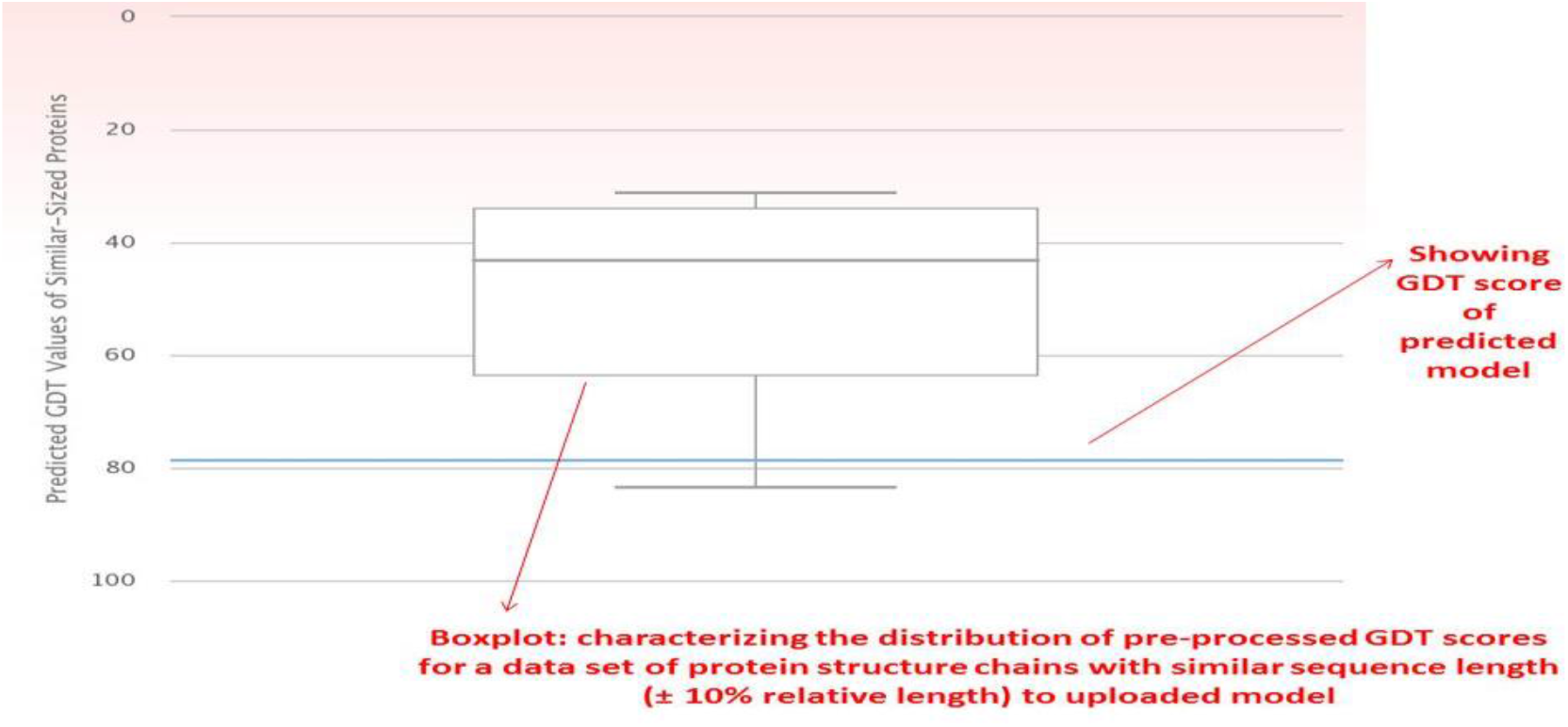
Global quality assessment of predicted model by eQuant.

**Fig. 12:**
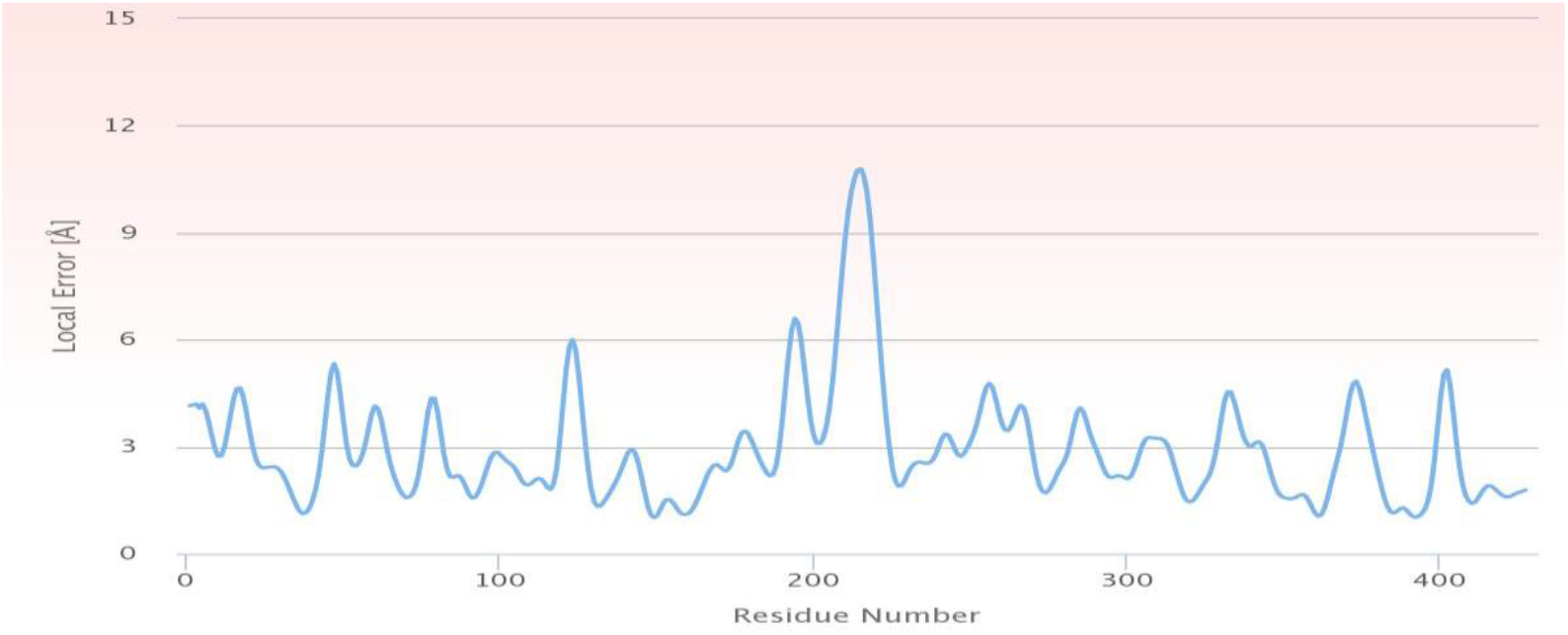
Local quality assessment of predicted model by eQuant.

**Fig. 13:**
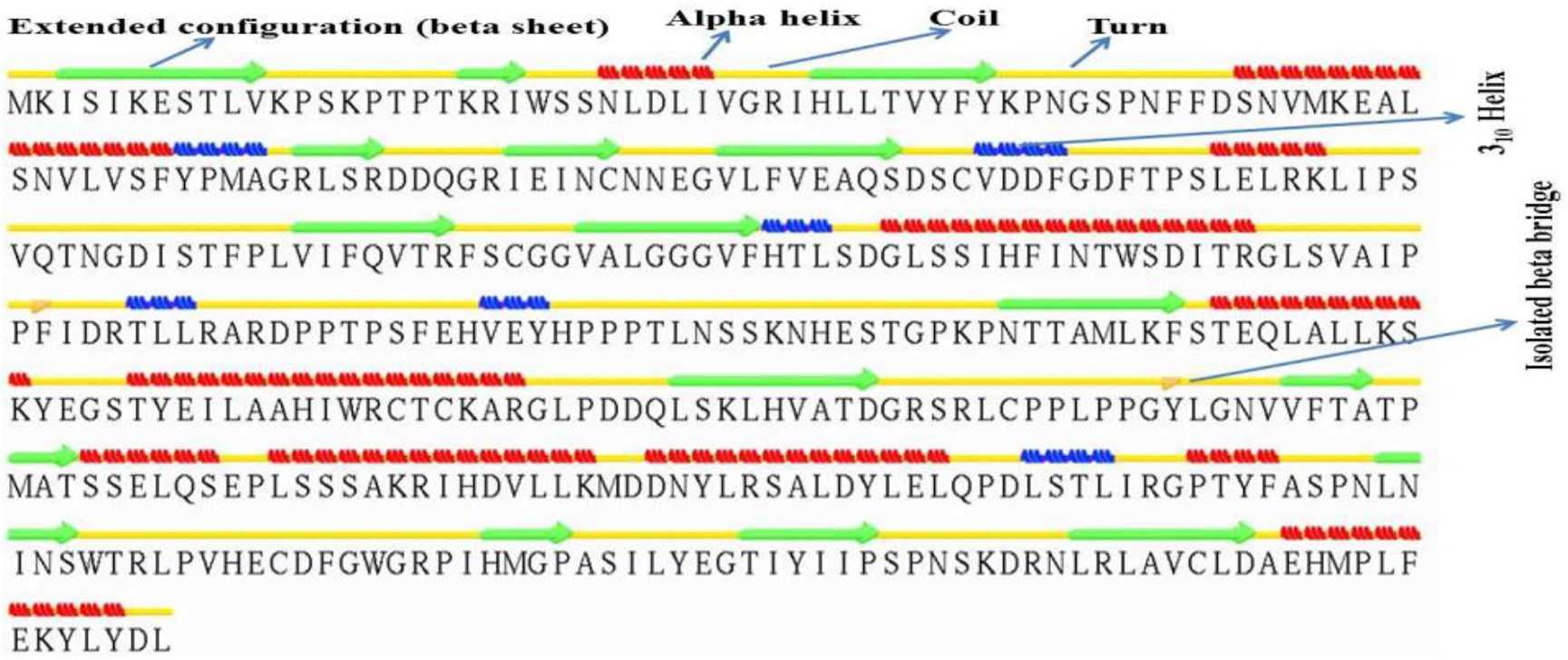
Secondary structure alignment of modeled SmHQT protein of *S. melongena.*

### Functional annotation

The predicted 3D structure of SmHQT of *S. melongena* and further analysis provided us a basis that helps to understand the biochemical function of the protein. 3D structure-based prediction revealed significant biochemical function and metabolic processes for SmHQT protein including catalytic activity, transferase activity etc. (Table 5).

**Table 5:** Predicted biological function of SmHQT using ProFunc server.

| Protein | Cellular component | Biological Process | Biochemical Function |
| --- | --- | --- | --- |
| SmHQT<br>ProFunc ID: bn90 | Cytoplasm, cell, cell part, intracellular | metabolic process, primary metabolic process, cellular process, cellular metabolic process | catalytic activity, transferase activity, transferase activity\, transferring acyl groups, transferase activity\, transferring acyl groups other than amino\acyl groups |

### Structure-based protein stability prediction

**Mutation sensitivity profile** was calculated for 3-D structure of SmHQT protein using MAESTRO web server. Colours represent the median value whereas sphere sizes the variance in ΔΔG. (Fig. 14). ΔΔGpred.<0.0 indicates a stabilising mutation.

**Fig. 14:**
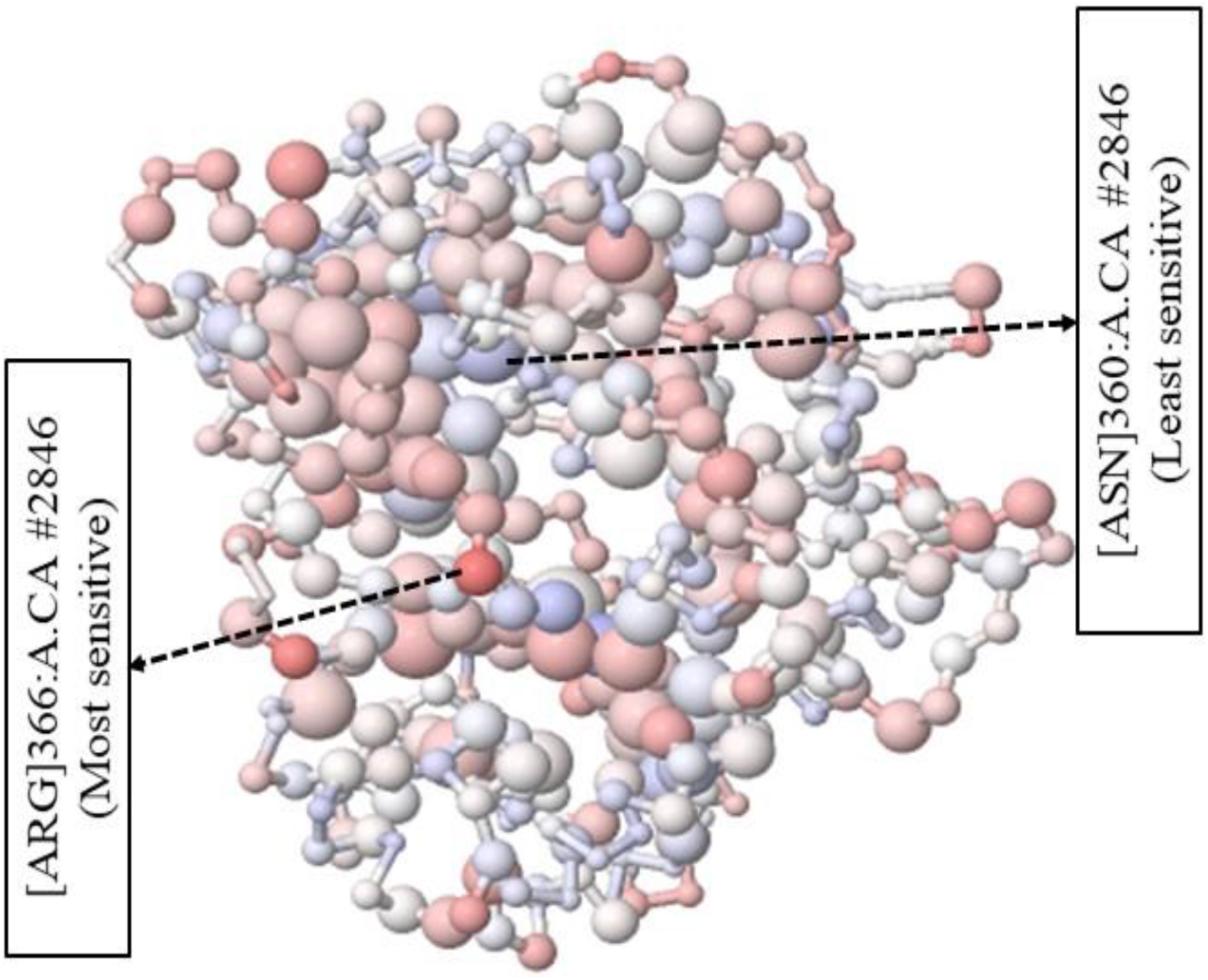
Diagrammatic representation of residues having different sensitivity to mutations.

Further, this web server provided a particular mode for the prediction of suitable disulfide bonds to stabilise a protein structure. Residue pairs with a Cβ-Cβ distance closer than 5Å are considered as potential binding partners. The results are sorted by bond score (Sss), from best (most negative score) (Fig. 15) to worst (highest value), which combined the predicted ΔΔG value and disulfide bond geometry penalties of residue pairs. Predicted potential disulfide bonds with their ΔΔGpred, Cpred and Sss scores are also given, where ΔΔGpred<0.0 kcal/mol indicates a stabilising mutation and Cpred is expressed as a value between 0.0 (not reliable) and 1.0 (highly reliable).

**Fig. 15:**
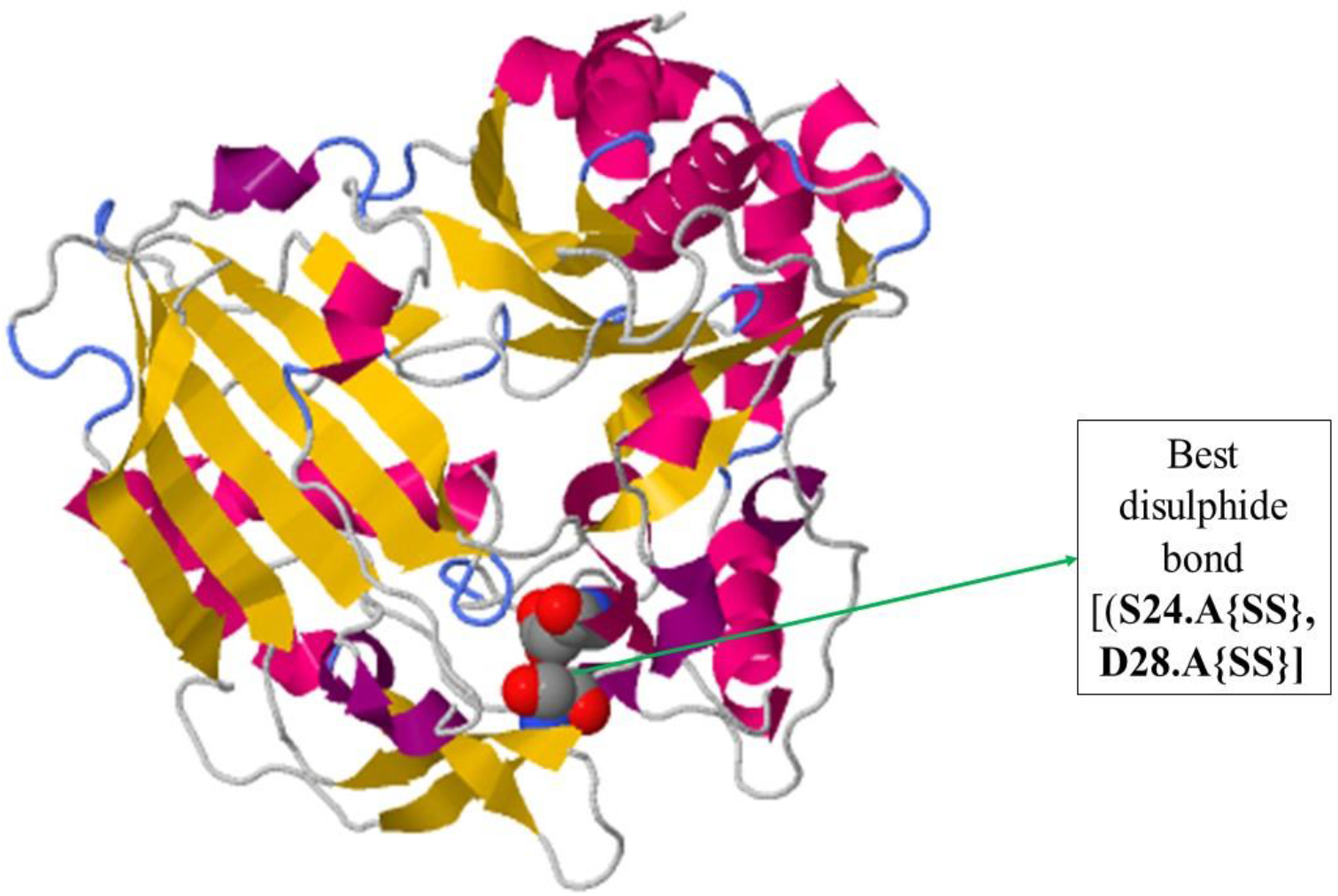
Structure of modelled protein showing best disulphide bond

### Detection of protein-protein interaction sites

The score of permanent/transient interaction ranges from 0.0 to 1.0 with 1.0 being the most confident score. For our protein, the overall score was 1.00 which suggests a transient type of interaction. SR and SR# stand for surface residues and residue number of surface residues, respectively; dL score is a likelihood of residue present at the interface; dL Z-score is the Z-score of dL score; tL-score is the likelihood score that a residue has permanent or transient interaction; tL Z-score is Z-score of tL score, where residues with a negative or positive value is predicted to have transient or permanent interaction, respectively. Residues at predicted protein binding site are highlighted in grey and residues predicted to have transient or permanent interaction are coloured in blue or red respectively (Fig. 16).

**Fig. 16:**
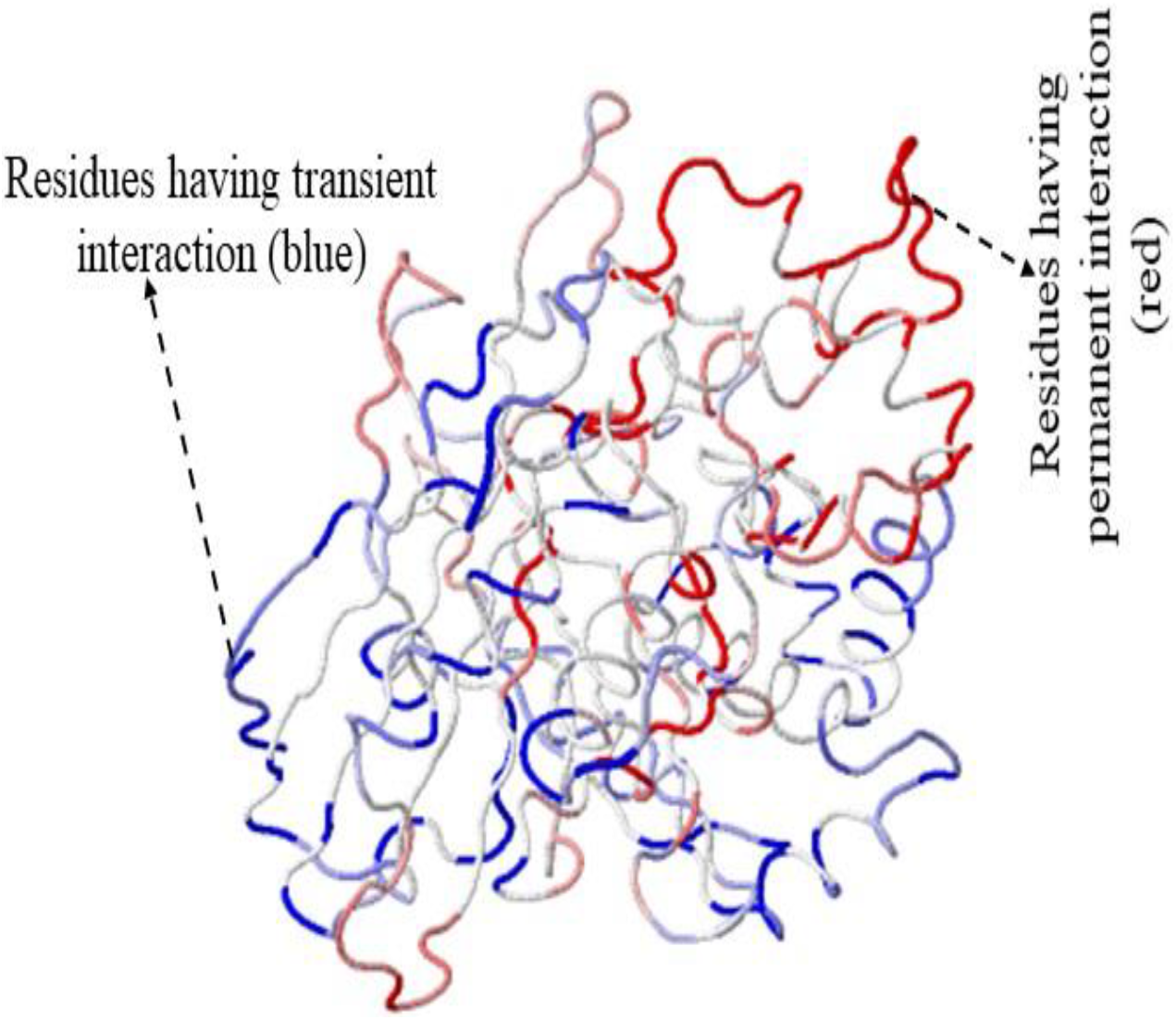
Visualization of interactive protein interface prediction

### Molecular Docking

Docking of SmHQT modelled protein (model002) with ligand quinic acid was carried out with SWISSDOCK web server based on EADock DSS (Grosdidier *et al.*, 2011) and total number binding clusters were also generated. From those clusters for model 002, cluster 10 was selected with the values for FullFitness to be −1951.9 kcal/mol and Estimated ΔG to be −7.87945 (kcal/mol) (Table 6). Protein-ligand binding for this cluster was visualised by using UCSF Chimera (Fig. 17).

**Fig. 17:**
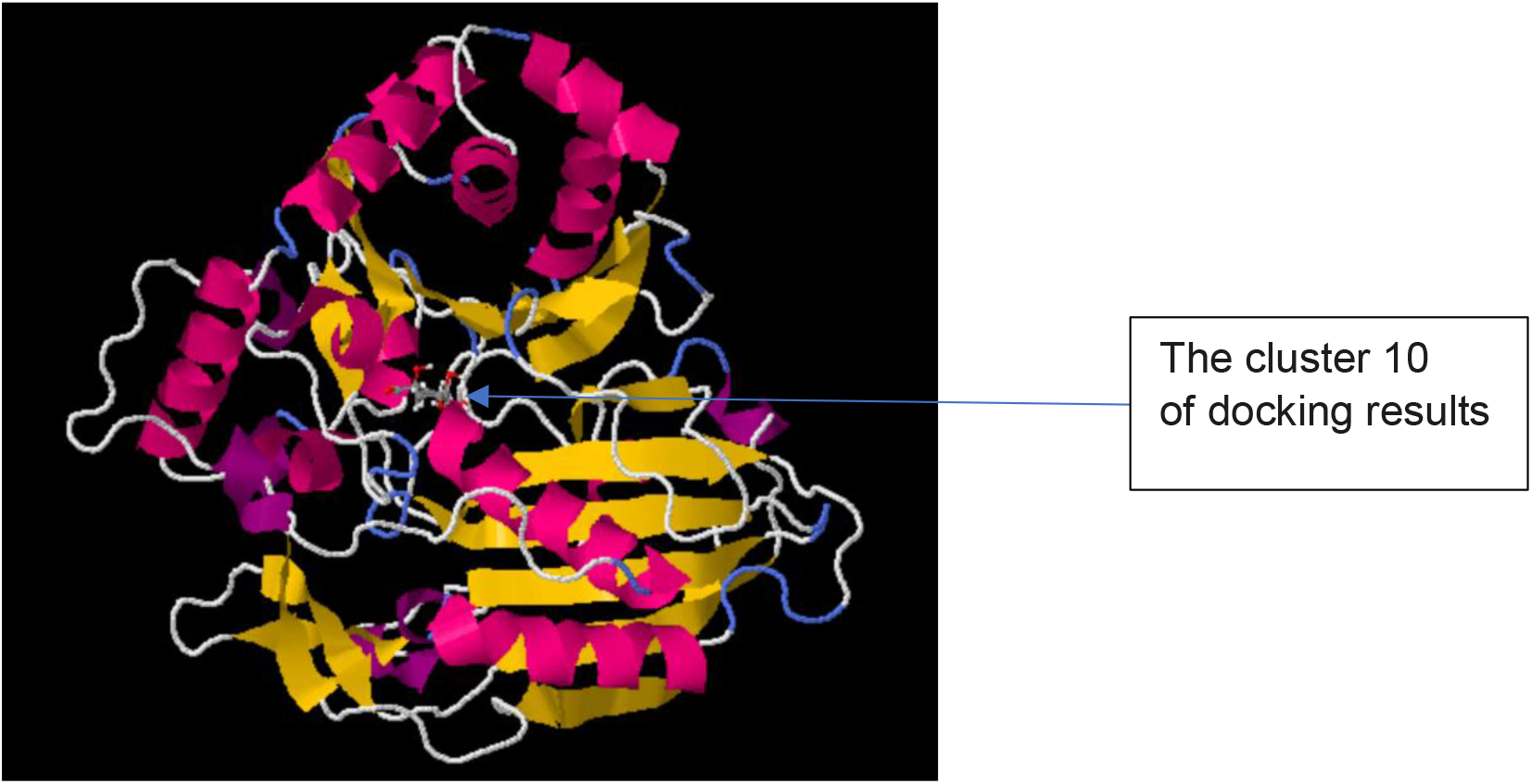
Ligand quinic acid binding within modelled protein SmHQT (model002).

**Table 6:**
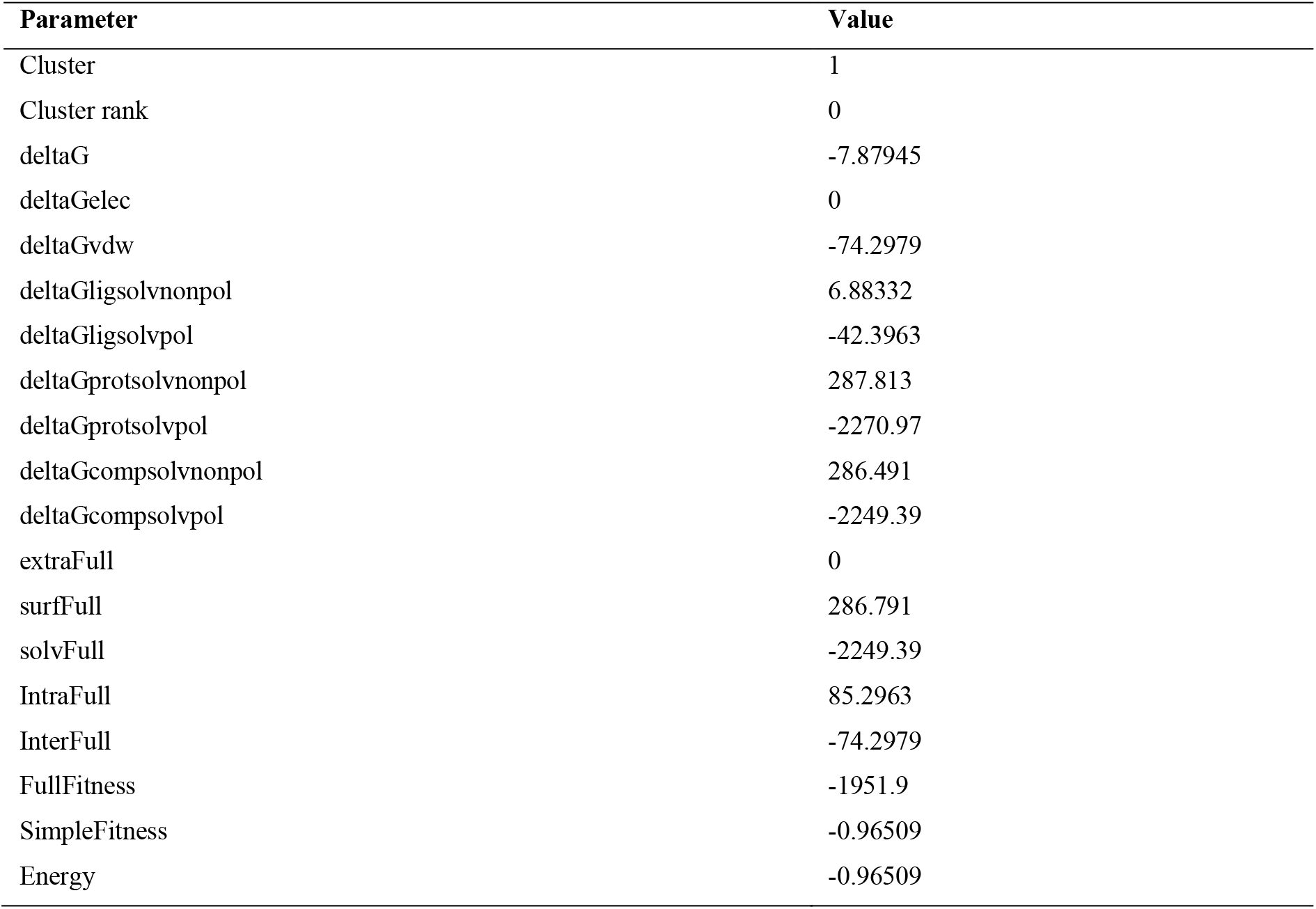
Docking results with SWISSDOCK.

## Discussion

Chlorogenic acid production in eggplant depends on the SmHQT protein. Even the eggplant genome has been sequenced. But there is not any detailed information regarding the SmHQT; therefore, here firstly we have analyzed the amino acid sequences present in the SmHQT of eggplant. After that, using homology modelling methods 3D model of SmHQT was predicated. Subsequently, several properties of modelled structure like stereochemical properties etc. were studied.

An attempt to generate a homology model of SmHQT protein was carried out using several tools. GO analysis suggested that SmHQT protein possess a transferase activity about its functions of transferring acyl groups (Kelley *et al.*, 2015; Schwede, 2013). The amino acid sequence was identified using HHsearch and BLAST. Top ten hits from each method were selected by coverage, lowest e-values, good resolutions, good identities and then used as templates for homology modelling (Gupta *et al.*, 2017). After that, ligands and ions were utilized for the modelling procedure. MODELLER 9v15 was used to create the homology model of SmHQT protein of *S. melogena.* A total of ten models were produced, and their discrete optimised potential energy (DOPE) scores were captured. The model002 having maximum score was deemed as the best model of SmHQT of S. melongena. PROCHECK, VERIFY3D and ERRAT programs authenticated the predicted model (Lahiri *et al.*, 2012). Ramachandran plot of the modelled protein presented the distribution patterns of residues in different regions. Three hundered forty residues out of total were in the favoured region of the Ramachandran plot (Hollingsworth and Karplus, 2010). Similarly, results obtained from PROCHECK indicated the stability of the selected protein model. The QMEAN Z-score of the model indicated a medium quality of protein model (Benkert *et al.*, 2011).

Whereas, the selected model appears to be of good quality, as inferred firstly from the appropriate values of QMEAN4 and secondly from the high values of quality factors determined by ERRAT and those of 3D-1D score estimated by VERIFY-3D. Global structure evaluation was also performed with PROSESS. Our predicted model accomplished overall quality, covalent bond quality, non-covalent/packing quality and torsion angle quality within an acceptable range (Sefid *et al.*, 2013). Assessment with eQuant predicted GDT, Z-score and TM-score provided a perfect match (Kryshtafovych, 2014). This prediction of change in stability upon point mutations in proteins may be valuable in further protein analysis and engineering. Detection of protein-protein interaction sites, prediction of protein-protein interaction sites in a protein structure may be exploited to get insights into the mechanism of protein function (Strokach *et al.*, 2019). Molecular docking has identified the algorithm that has pulled the structure inside (Pettersen *et al.*, 2004).

Overall, we analysed the sequence of the SmHQT protein of eggplant, thereby, defining various essential functions that this protein might perform. Further, homology modelling provided its detailed 3D structure model. The model developed has a good overall parameters for its structure related aspects. Moreover, we have validated it using several programs including PROCHECK. Molecular docking was also performed in SwissDock using ligand quinic acid, and a cluster was selected based on the value of FullFitness.

