## Supplementary Fig. 1,2,3 for "Sequence Analysis and Homology Modelling of SmHQT Protein, a Key Player in Chlorogenic Acid Pathway of Eggplant"

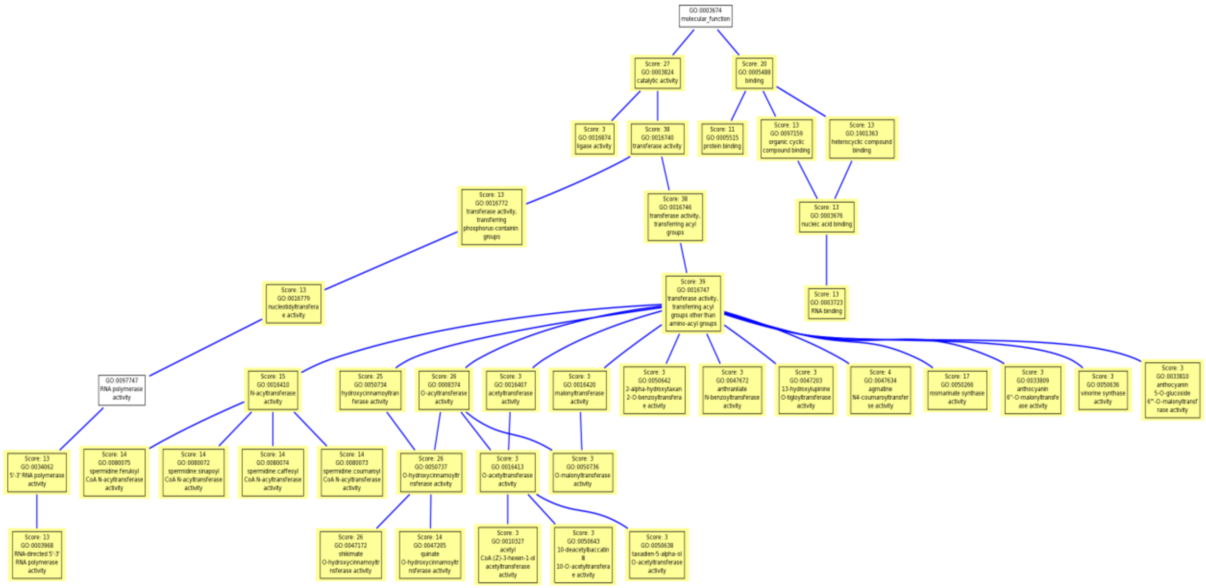


**Supplementary Fig. 1:** Molecular function ontology.

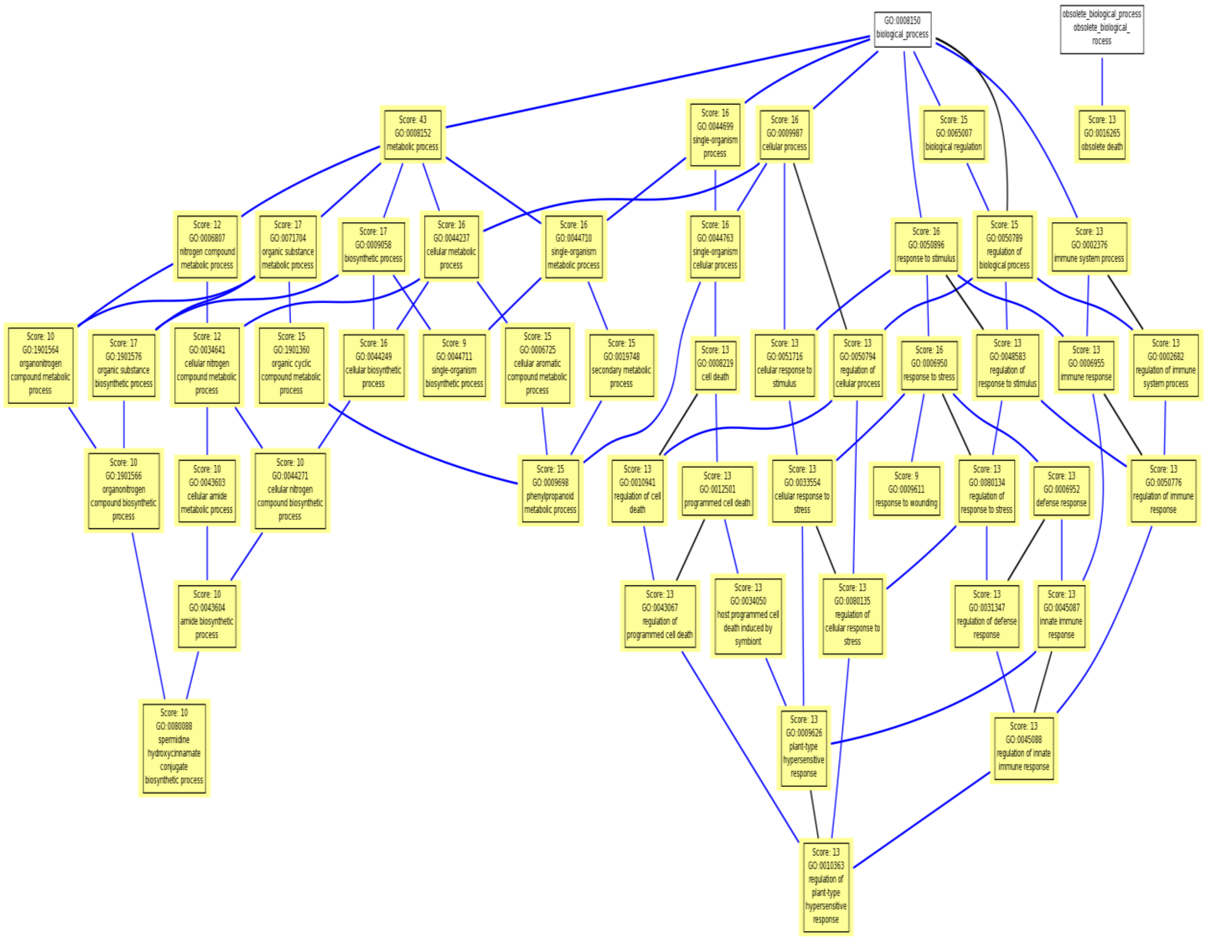


**Supplementary Fig. 2:** Biological process ontology.

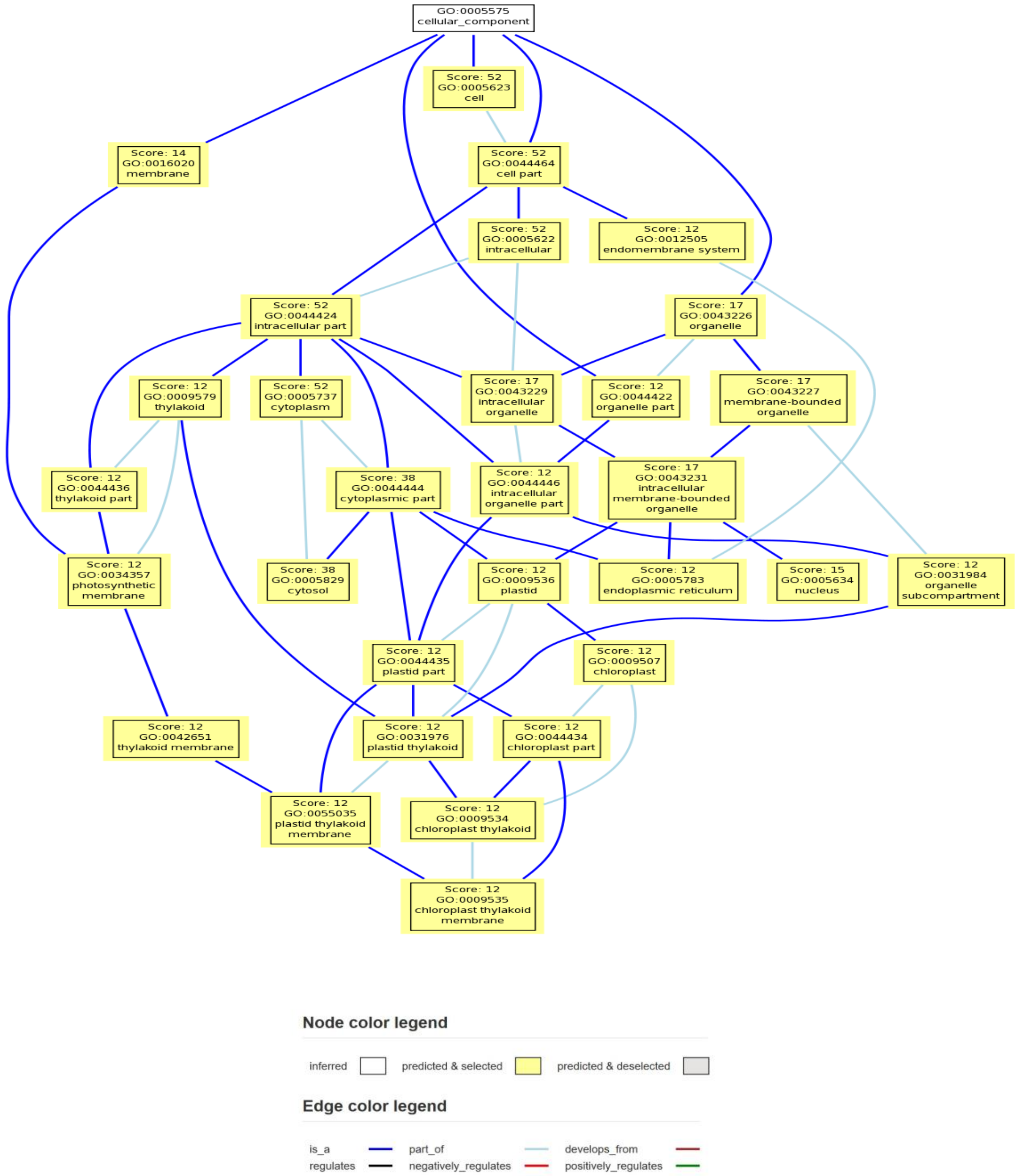


**Supplementary Fig. 3:** Cellular component ontology.

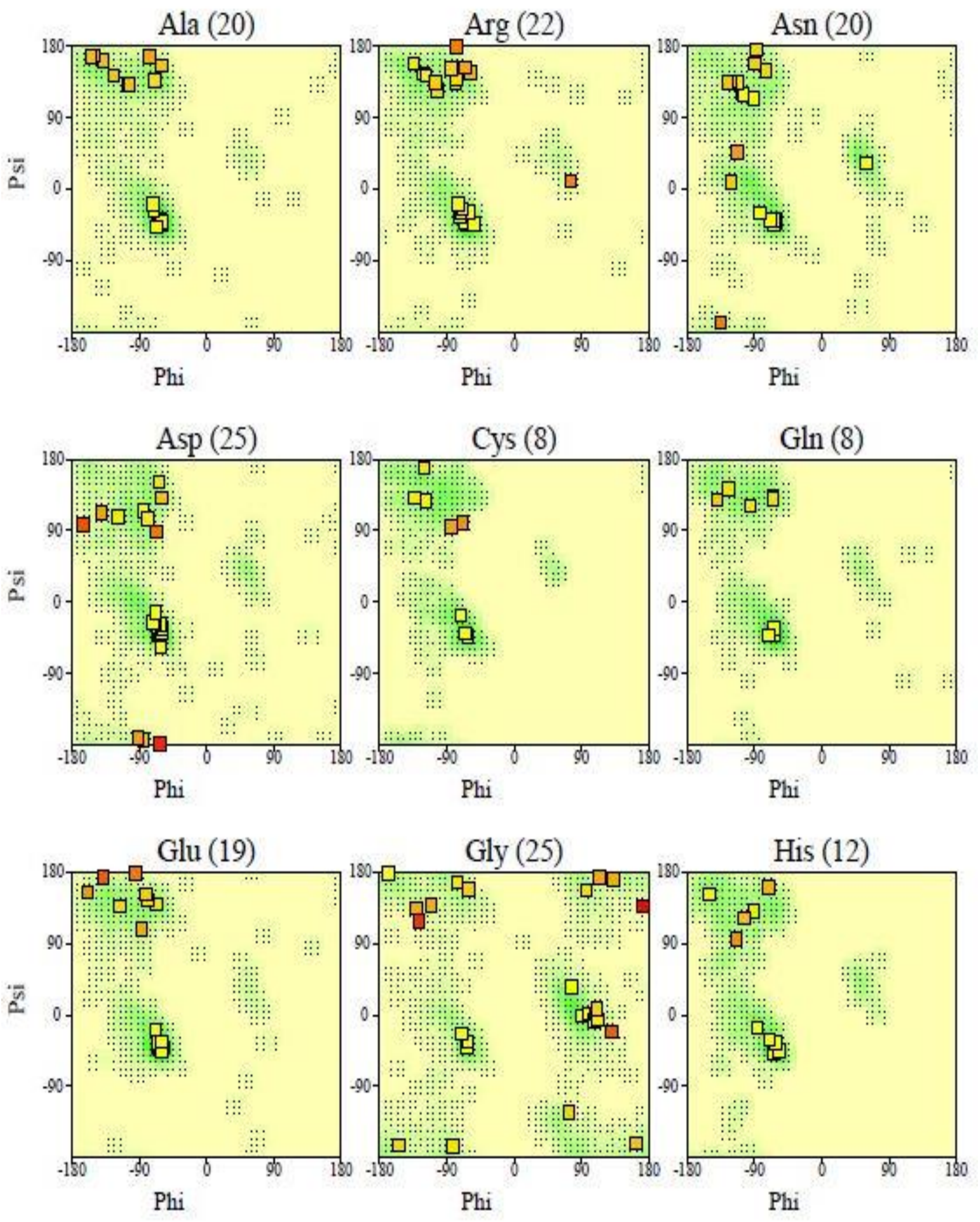


Numbers of residues are shown in brackets. Those in unfavourable conformations (score < -3.00) are labelled.
Shading shows favourable conformations as obtained from an analysis of 163 structures at resolution 2.0A or better.

**Supplementary Fig. 4:** Figure showing separate Ramachandran plots for each of the 20 different amino acid types.

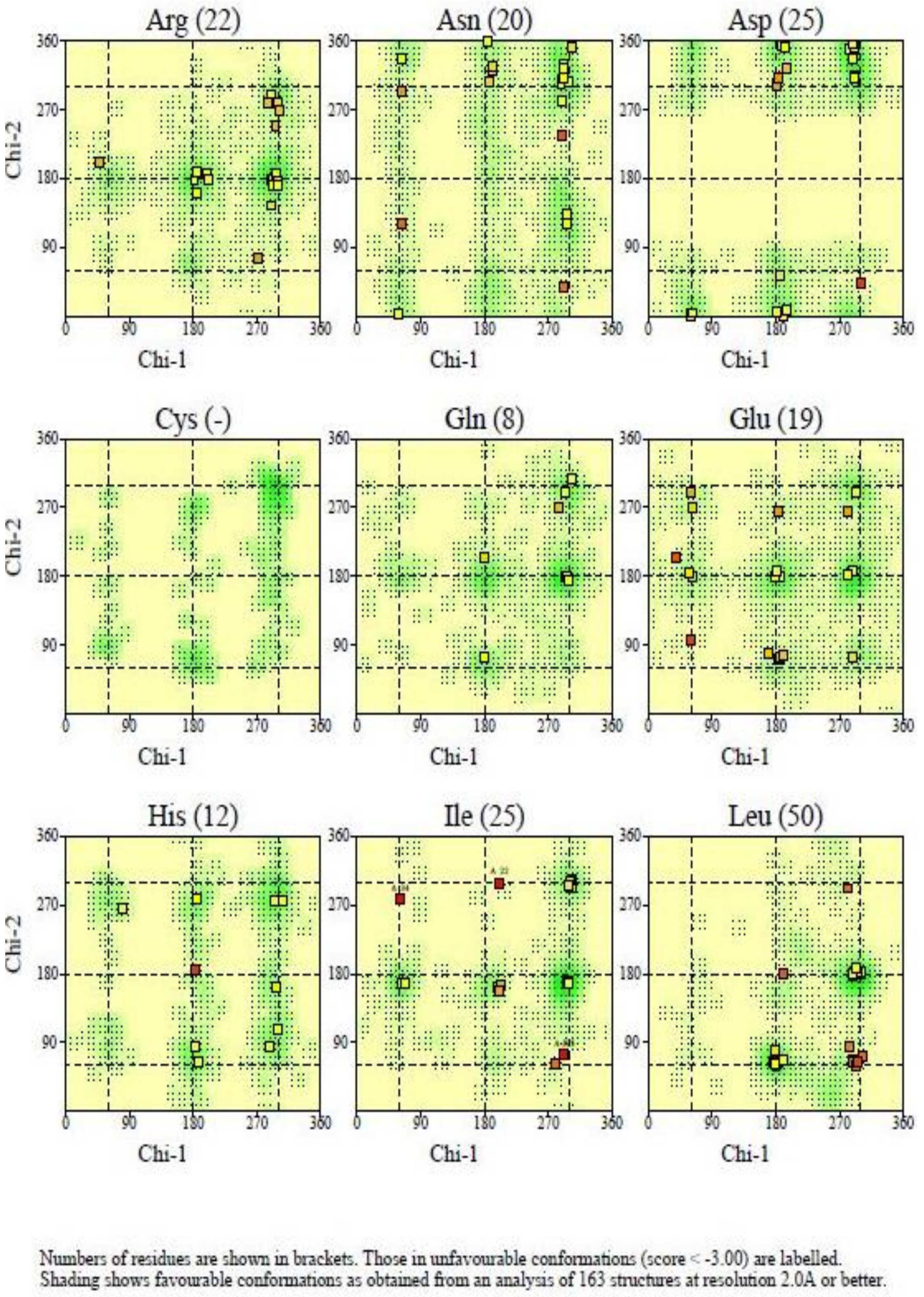


**Supplementary Fig. 5:** Main-chain bond lengths (dark shading on Chi1-Chi2 plots indicating favourable regions covered by residues on plots).

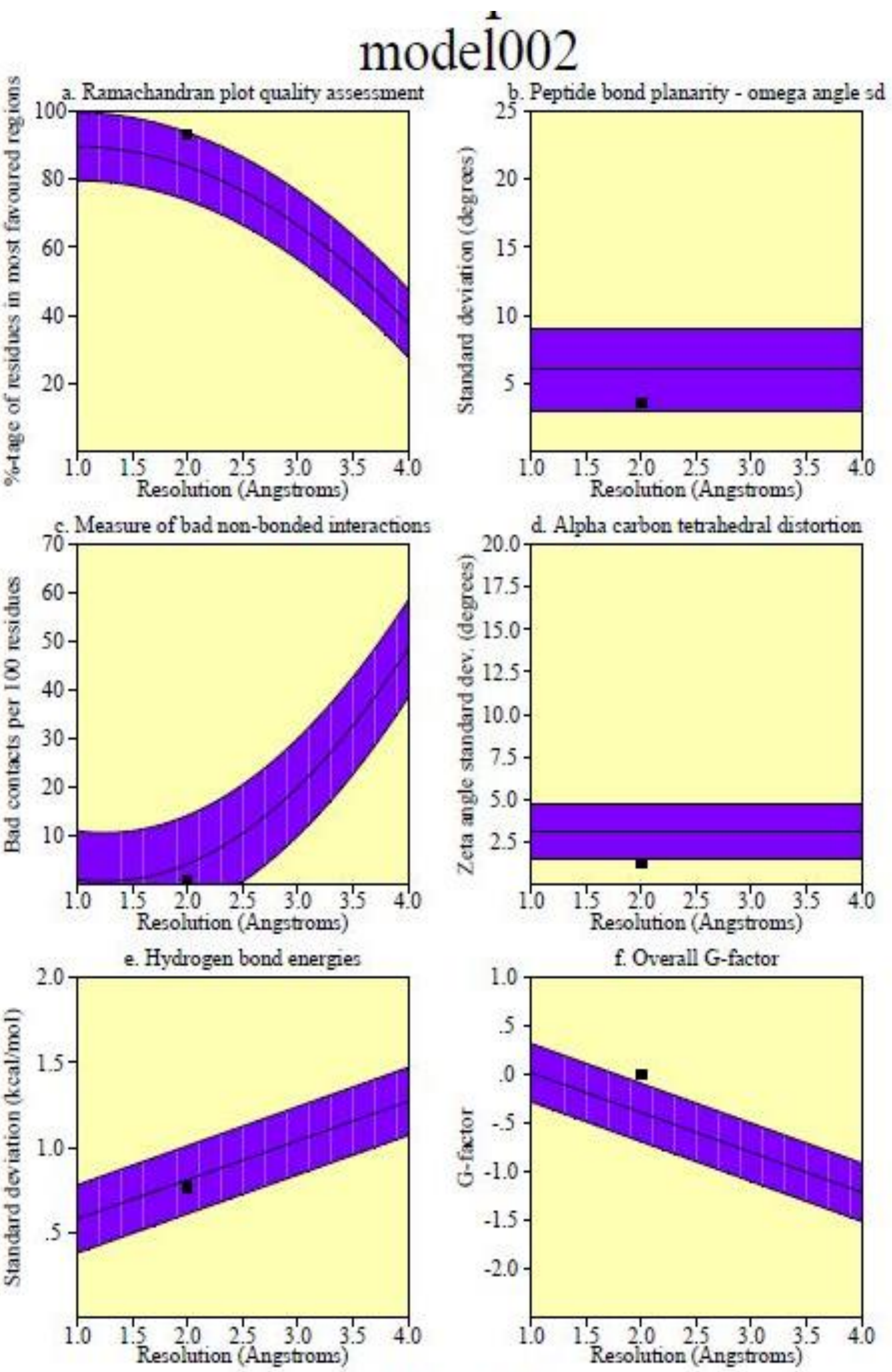


Plot statistics

| Stereochemical parameter | No. of data pts | Parameter value | Comparison values Typical value | Band width | No. of band widths from mean | |
|---|---|---|---|---|---|---|
| a. %-tage residues in A, B, L | 365 | 93.2 | 83.8 | 10.0 | .9 | Inside |
| b. Omega angle st dev | 424 | 3.5 | 6.0 | 3.0 | -.8 | Inside |
| c. Bad contacts / 100 residues | 4 | .9 | 4.2 | 10.0 | -.3 | Inside |
| d. Zeta angle st dev | 402 | 1.3 | 3.1 | 1.6 | -1.2 | BETTER |
| e. H-bond energy st dev | 248 | .8 | .8 | .2 | -.2 | Inside |
| f. Overall G-factor | 427 | .0 | -.4 | .3 | 1.3 | BETTER |

**Supplementary Fig. 6:** Chi1-Chi2 plots (showing the values of all six parameters).

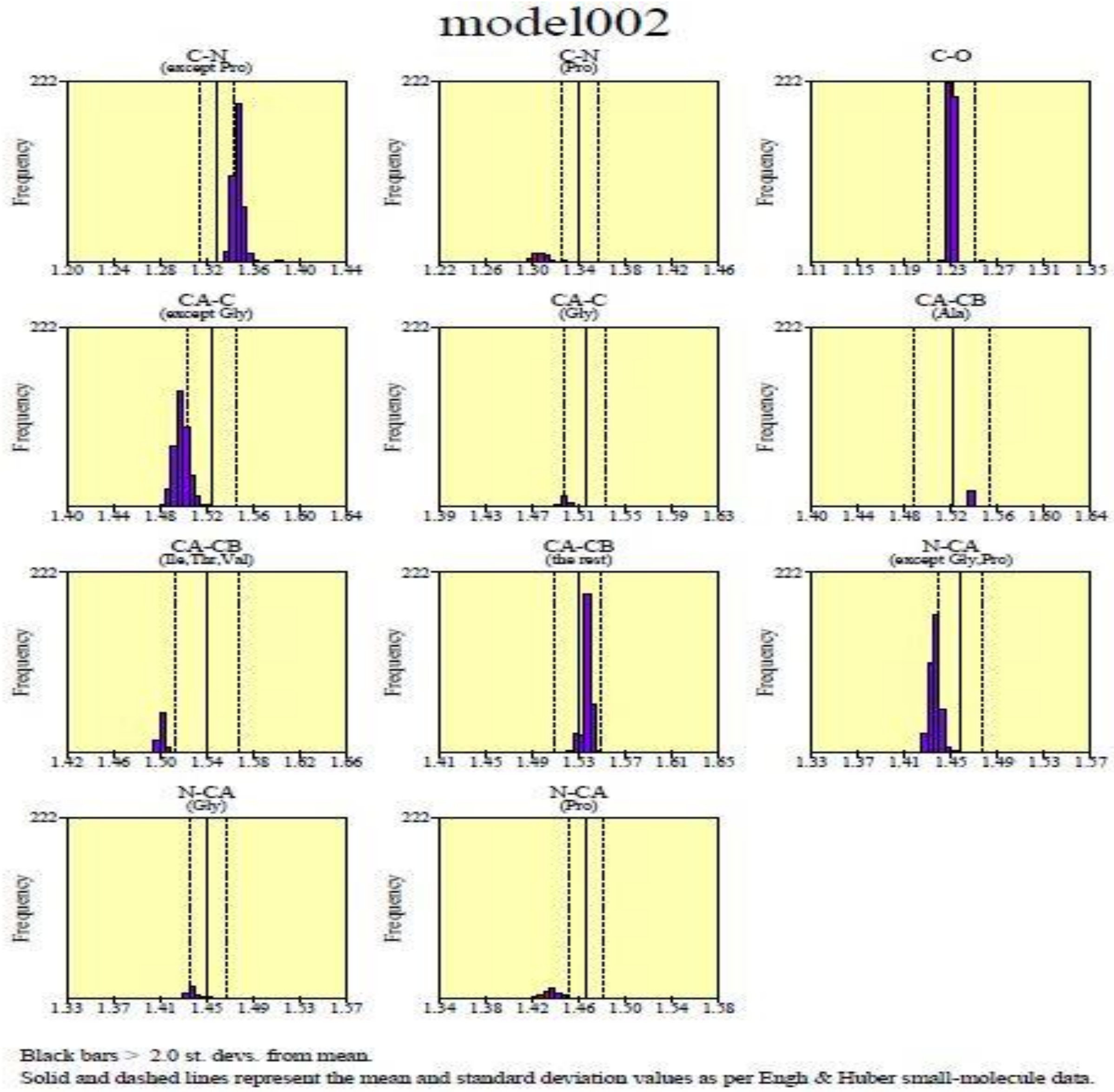


**Supplementary Fig. 7**: Main-chain bond lengths (showing the distributions of each of the different main-chain bond lengths in the structure)

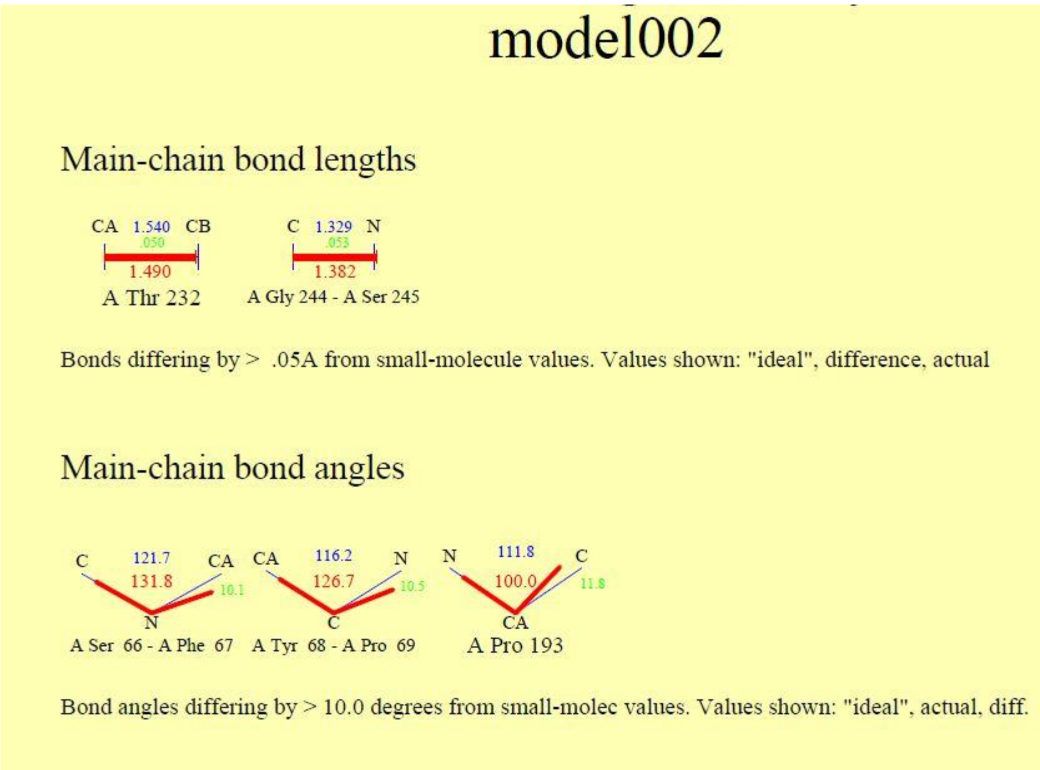


**Supplementary Fig. 8:** Distorted geometry
